# Neural correlates of kinematic features of passive finger movement revealed by univariate and multivariate fMRI analyses

**DOI:** 10.1101/2024.11.20.624514

**Authors:** Gustavo S. P. Pamplona, James Sulzer, Ewa Beldzik, Olivier Lambercy, Silvio Ionta, Roger Gassert, Jarrod Lewis-Peacock

## Abstract

Finger movements are associated with a relatively large neural representation. Passive finger movement – which involves refraining from actively performing or resisting movement – is a robust approach to investigate the neural representation of kinesthesia and proprioception in the brain. While some studies have characterized the neural correlates of passive finger movement, they have relied solely on mass univariate analysis, potentially affecting result sensitivity. Additionally, limited consideration has been given to stimulus duration, a factor closely tied to some kinematic features (amplitude and velocity), which recently proposed modeling approaches now take into account. Here, we reanalyzed previously published data using univariate and multivariate analysis to understand how kinesthesia is neurally encoded in neurotypical subjects in two separate experiments. Systematic passive stimulation of the fingers was provided using an MR-compatible robot while functional magnetic resonance imaging data was recorded. Our analyses consisted of univariate and multivariate approaches, conducted separately for each kinematic feature and adjusted for stimulus duration, regardless of whether brain activation scales with it. We provide a detailed mapping of brain areas related to amplitude, velocity, and direction of passive finger movement, including sensorimotor, subcortical, and cerebellar areas. In general, multivariate pattern analysis was more sensitive than the univariate approach in identifying brain regions associated with passive finger movement. Our univariate analysis demonstrated that activity in sensorimotor and subcortical areas was higher for larger amplitudes and slower velocities, which opposes to the original study’s results, likely due to our treatment of stimulus duration as a confounder specified as a parametric modulator. A novel result, we also demonstrated that brain activity in sensorimotor areas was higher for extension compared to flexion of passive finger movement. In terms of kinematic features, a larger neural representation was found for amplitude and direction compared to velocity of passive finger movement. This indicates that kinesthesia and proprioception may be more reliant on displacement than kinematic aspects of passive finger movement. While univariate analyses are limited in addressing spatial heterogeneity and subject-level variability, our multivariate analyses showed increased sensitivity in identifying brain regions encoding passive movement. Our findings may extend the knowledge of how the brain encodes physical movements and may help design neurorehabilitation strategies.

## 1. Introduction

Passive movement entails moving the body part of an individual, who refrains from performing active movement and from resisting against an external source of movement. At a neuroscientific level, passive movement of body parts can be employed to study kinesthesia and proprioceptive processes in the brain and to consolidate neuroscientific theories of motor control. The brain correlates of passive movement have been studied extensively using various neuroimaging techniques for several body parts, such as the wrist (Bae et al., 2017), hand (Boscolo Galazzo et al., 2014; Guzzetta et al., 2007; Zhavoronkova et al., 2017), arm (Estévez et al., 2014; Weiller et al., 1996), lower limb (Jaeger et al., 2014), and finger (Dueñas et al., 2018; Lolli et al., 2021; Mima et al., 1999; Nurmi et al., 2018; Sugawara et al., 2016). Hand and fingers have a proportionally large somatotopic representation compared to other body parts (Penfield and Rasmussen, 1950). Passive finger movement commonly elicits responses in the supplementary motor area (SMA), the contralateral postcentral (primary somatosensory cortex – S1) and precentral gyrus (primary motor cortex – M1), and the bilateral secondary somatosensory cortex (S2), thalamus, putamen, and ipsilateral cerebellum (Dueñas et al., 2018; Lolli et al., 2021; Mima et al., 1999; Nurmi et al., 2018; Sugawara et al., 2016). This pattern for passive movement is similar to the one associated with active movement (Boscolo Galazzo et al., 2014; Guzzetta et al., 2007; Weiller et al., 1996), but with a relatively weaker strength of brain activity (Lolli et al., 2019; Mima et al., 1999; Zhavoronkova et al., 2017). Early work on peripheral receptors showed that kinematic features of passive movement are dissociated from each other (McCloskey, 1973; Proske and Gandevia, 2009; Sittig et al., 1985), suggesting that kinematic features (such as amplitude, velocity, and direction) could also be represented differentially in the brain.

Proprioception is especially important for the dexterous movements of the hand and fingers, as it enables precise control and awareness of position without visual guidance. Robotically-driven motion, compared to manual stimulation by an experimenter, offers a systematic way to provide subjects with reproducible and stereotyped passive movement of body parts (Aggogeri et al., 2019). When robotics is used in conjunction with functional magnetic resonance imaging (fMRI) through an adapted and compatible construction (Gassert et al., 2006), one can study the passive finger movement using high-resolution imaging and well-controlled kinesthetic stimulation (Dueñas et al., 2018). However, despite the high spatial resolution, the characterization of passive finger movement in the brain has only been studied using mass univariate analyses (Boscolo Galazzo et al., 2014; Dueñas et al., 2018; Lolli et al., 2021, 2019; Nurmi et al., 2018), i.e., obtaining large clusters of spatially convergent voxels across subjects with concordant activity magnitude.

Furthermore, mass univariate analyses may be insensitive to patterns of activity across spatially distributed voxels (Davis et al., 2014), i.e., their results depend on how spatially homogeneous the activation is. Alternatively, multivariate pattern analysis (MVPA) applied to fMRI offers greater sensitivity for characterizing the neural underpinnings of a given task, stimulation, or behavior (Haxby et al., 2014; Hebart et al., 2015; Kriegeskorte et al., 2006; Norman et al., 2006). Such an approach uses machine learning methods to discriminate between conditions that are reliably associated with more distributed encoded brain patterns (Averbeck et al., 2006; Kriegeskorte et al., 2006; Riggall and Postle, 2012). Compared to univariate analysis, MVPA can detect spatial heterogeneity and neglects subject-level variability (Coutanche, 2013; Davis et al., 2014). Consequently, MVPA is powerful to differentiate between conditions that would be intertwined with traditional univariate approaches and generate spatially accurate brain maps that extend our understanding of neural functioning across subtle experimental variations.

Employing and comparing univariate and multivariate methods, as well as modeling inherent confounds, may substantially advance our comprehension of brain correlates of passive finger movement. Here we performed new analysis on a previously recorded dataset (Dueñas et al., 2018). This dataset comprised of high-resolution fMRI data from 41 healthy subjects in two independent experiments while an MRI-compatible robotic system provided finger displacement according to different kinematic features. Using univariate and multivariate techniques, we identified neural responses associated with three different kinematic features (amplitude, velocity, and direction) of passive finger movement. We hypothesized that the univariate and multivariate representations would exhibit heightened sensitivity to amplitude of passive finger movement compared to other kinematic features, reflecting brain responses that rely primarily on the proprioceptive input to interpret passive movement. This hypothesis is based on previous findings showing that sensorimotor activity (measured by a univariate analysis and controlled by stimulus duration) due to active finger movements is more specific to amplitude than other kinematic features (Shirinbayan et al., 2019). By providing linear (and not oscillatory) movements experimentally and by using a model that controls for false positive rates regardless of whether brain activation scales with stimulus duration, we provide unique insights on the neural responses to passive finger movement. Our directional results of univariate analysis and improved sensitivity of multivariate analyses deepen our understanding of how neural correlates of passive finger movement are characterized.

## 2. Methods

### 2.1. Participants

We performed new analyses of a previously recorded dataset (Dueñas et al., 2018). The data consisted of anatomical and functional MR images collected at the MRI Center of the Psychiatric Hospital of Zurich, University of Zurich, Switzerland. Data were obtained from 41 right-handed healthy subjects who participated in two experiments, hereafter referred to as Exp 1 and Exp 2 (following the nomenclature of the original publication) (Dueñas et al., 2018). The dataset used for the present study contained only subjects whose head displacement during functional runs was no greater than 3 mm, filtered out in the original study. Nineteen subjects participated in Exp 1 (14 females, 27.8 ± 8.9 years) and 22 subjects (11 females, 25.2 ± 2.9 years) participated in Exp 2. The mean age of the participants was not different between the two experiments (two-sample t-test, p = 0.20). All subjects read and gave written consent for data reuse and received monetary compensation for their participation. The study was approved by the local ethics committee (KEK 2010-0190).

### 2.2. Experimental design

We acquired fMRI of the subjects’ brains while they were undergoing robotically-driven passive finger movement, provided through an MRI-compatible robotic system (Fig. 1A). Prior to the MRI recording, we measured the maximum aperture between right thumb and index finger for each subject with a ruler outside the scanner room. Subjects were laid on the scanner table in a supine position with the wrist in the anatomically neutral position. We attached the distal phalanges of the subjects’ right thumb and index finger using Velcro straps to the slave part of the robot (MRI-compatible), previously mounted on the right side of the scanner table. This part of the robot was controlled by the master part (non-MRI-compatible), lying outside the scanner room, and connected to each other through a hydraulic transmission. The robot was designed provide individuals with flexion and extension of the index finger relative to the thumb. The motion of the index finger was controlled during the experiments through an optical encoder. The robotic system has been validated against artifacts for concomitant fMRI recording and poses no safety concerns for participants (Gassert et al., 2006). A dedicated computer with custom code written in LabVIEW (Laboratory Virtual Instrument Engineering Workbench, National Instruments) controlled the robot. Subjects were familiarized with the experimental procedure before being positioned in the MRI scanner. They were instructed to remain relaxed, to not exert active movement in reaction to passive movement, to keep their eyes open, and to not fall asleep during the measurement.

**Figure 1.**
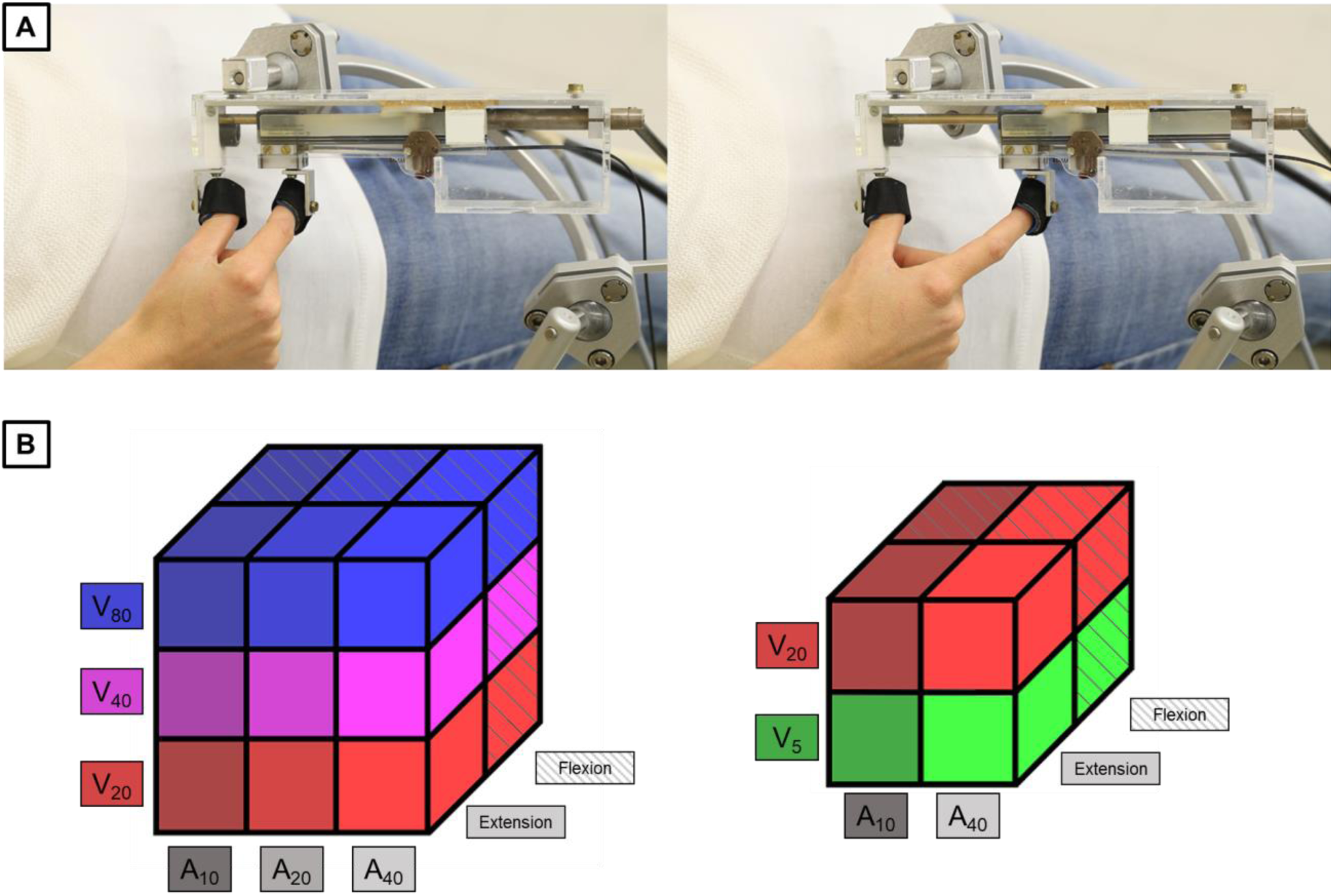
(A) MRI-compatible robot used in Exps 1 and 2. Adduction and abduction of right thumb and index finger relative to each other were provided to the subjects, while they were lying in the MRI scanner, with the wrist positioned in the anatomically neutral position. We show two static positions of the fingers between event-related trials. Relative to a null aperture (Velcro straps touching each other), the left/right image shows an aperture of 20%/80% of the subject’s maximum amplitude, respectively. (B) Factorial designs for Exps 1 and 2, respectively. Each element of the 3-D representation is a possible combination of levels of kinematic features (amplitude, velocity, and direction) in an event-related trial. Exp 1 was designed as a 3×3×2 factorial design, with three levels of amplitude, three levels of velocity, and two levels of direction. Exp 2 was designed as a 2×2×2 factorial design, with two levels of amplitude, two levels of velocity, and two levels of direction. Shading represents different levels of amplitude, colors represent different levels of velocity, and textures represent different levels of direction.

Each trial (robotically-driven passive movement) consisted of a single displacement of right thumb and index finger relative to each other for a combination of levels of kinematic features – amplitude, velocity, and direction (Fig. 1B). The order of the trials and the interval between trials were pseudorandomized for each subject prior to scanning. For Exp 1, trials consisted of a 3 x 3 x 2 combination of amplitudes of 10, 20, or 40% of the maximum aperture (A_10_, A_20_, and A_40_), velocities of 20, 40, or 80% of the maximum aperture per second (V_20_, V_40_, and V_80_), and directions of extension or flexion. The interval between trials was 4 ± 2 s for Exp 1, which constituted a fast event-related design. Each unique combination was repeated eight times per run in Exp 1. The run length was 13.5 min for Exp 1. The mean ± standard deviation of the amplitudes in Exp 1 across individuals was 9.6 ± 1.3 mm for A_10_, 18.1 ± 2.6 mm for A_20_, and 37.0 ± 5.3 mm for A_40_. The mean ± standard deviation of the (average) velocities in Exp 1 across individuals was 26.4 ± 3 mm/s for V_20_, 56.7 ± 7.1 mm/s for V_40_, and 134.6 ± 17.0 mm/s for V_80_.

For Exp 2, trials consisted of a 2 x 2 x 2 combination of amplitudes of 10 or 40% (A_10_ and A_40_) of the maximum aperture, velocities of 5 or 20% of a maximum aperture per second (V_5_ and V_20_), and directions of extension or flexion. The interval between trials was 12 ± 2 s for Exp 2, which constituted a slow event-related design. Each unique combination was repeated five times per run in Exp 2. The run length was 10.5 min for Exp 2. The mean ± standard deviation of the amplitudes in Exp 2 across individuals was 7.3 ± 1.3 mm for A_10_, and 38.3 ± 4.6 mm for A_40_. The mean ± standard deviation for the (average) velocities in Exp 2 across individuals was 6.8 ± 0.8 mm/s for V_5_, and 30.6 ± 2.6 mm/s for V_20_.

### 2.3. MRI acquisition

Imaging data were acquired with a Philips Achieva 3 T MR scanner and a 32-channel head coil. For both experiments, we first acquired functional volumes covering the whole cerebrum and the superior part of the cerebellum, obtained with a gradient-echo planar imaging sequence, 35 axial slices, repetition time/echo time (TR/TE) = 2000/35 ms, matrix size = 240 × 240 mm^2^, voxel size = 3 x 3 x 4.07 mm^3^, 1.1 mm slice gap, and 400 and 306 volumes for Exps 1 and 2, respectively. Next, we acquired high-resolution three-dimensional anatomical images covering the whole brain, obtained with a T1-weighted sequence, 180 sagittal slices, TR/TE = 6.9/3.1 ms, flip angle = 8°, matrix size = 256 × 256 mm^2^, voxel size = 1 mm^3^). The total duration of a session was about 60 min for each experiment.

### 2.4. Data analysis

#### 2.4.1. MRI preprocessing

All MRI data acquired during passive finger movements were preprocessed prior to statistical mapping analysis, using SPM (Statistical Parametric Mapping, Wellcome Trust Centre for Neuroimaging, www.fil.ion.ucl.ac.uk/spm/) and custom scripts written in MATLAB (Mathworks, Natick, MA). We first slice-time corrected the functional images for each run and subject. We then estimated the six translation and rotation parameters of head motion and used them to spatially realign the functional images to a mean referential image. Next, we coregistered the anatomical image to the mean functional image. Thus, we used the coregistered anatomical image to create a deformation field, which was used to normalize all functional images to the Montreal Neurological Institute (MNI) space. In this process, we resampled the voxel size to 3 mm^3^. Finally, functional images were spatially smoothed using an isotropic Gaussian kernel of 6 mm^3^ full-width at half maximum, to be used only for univariate analysis.

#### 2.4.2. Estimation of activation for factorial design

We estimated voxel-wise activation for each unique combination of kinematic feature levels using a general linear model (GLM) for each subject. The estimation process was performed twice: once with unsmoothed functional MR images for MVPA, and once with smoothed images for univariate analysis. For each run, we designed a matrix with eighteen regressors (3 amplitude x 3 velocity x 2 direction levels) for Exp 1 and eight regressors (2 amplitude x 2 velocity x 2 direction levels) for Exp 2. Depending on the scientific question, time-on-task effects can act as confounding factors in univariate analysis (Domagalik et al., 2014; Yarkoni et al., 2009). Additionally, stimulus duration can confound MVPA (Todd et al., 2013), potentially causing the classifier to be trained on stimulus duration rather than the condition of interest per se. To account for this confound, we adjusted the activation estimation in run-specific regressors by treating stimulus duration as a covariate of no-interest. Accounting for trial-specific time-on-task differences in the model was first proposed by Grinband et al. (2008), in which the trials were modeled as boxcar functions with varying durations, according to the trial – the so-called variable epoch model. This modeling approach was initially applied in the study by Dueñas et al. (2018), which first described the data used in our present study. However, recent methodological work demonstrates that, while the variable epoch model effectively controls for false positives in cases where neural activity duration scales with time-on-task, it leads to a high rate of false positives when neural activity duration does not scale with time-on task (Mumford et al., 2024). Therefore, a model that minimizes false positives in both scenarios – whether neural activity duration scales or does not scale with time-on-task – is desired. Fulfilling these requirements, a proposed solution is a model uses constant regressors modulated by stimulus duration (Mumford et al., 2024). We adopted this approach to account for stimulus duration to keep the rate of false positives low, regardless of whether activation correlates with stimulus duration. Furthermore, we assumed a full interaction model, in which we accounted for neural activation that could vary with stimulus durations at different levels for each condition (Mumford et al., 2024). Therefore, we modulated each condition-specific constant regressor separately by stimulus duration. These constant regressors were constructed using boxcar functions, based on the onset of each robotic displacement and the mean stimulus duration across all trials within a run for each subject. For each constant regressor, a parametric modulator was added by specifying the mean-centered stimulus duration for each trial within each run. All parametric modulators were modeled as first-order (i.e., linearly associated). Additional regressors representing the covariates of no-interest, namely six head motion parameters and the runs, completed the design matrix. Each regressor was convolved with a canonical hemodynamic response function provided in SPM. We then simultaneously estimated voxel-wise the subject-specific GLM betas for each regressor. For each GLM, simple contrast maps of interest were created using the beta estimates for the constant regressors representing the unique combinations of kinematic feature levels. We used a high-pass filter of 128 s, a liberal masking threshold of 0.1, and an auto-regressive model of first degree to remove autocorrelation in the signal.

As described, we assumed that BOLD signals could potentially relate to stimulus durations at different levels for each condition in our model, which constitutes a full interaction model (Mumford et al., 2024). Therefore, modulated regressors were added to each condition-specific regressor. However, one could also assume that BOLD signals would relate to stimulus durations equally across conditions, without an interaction effect. To explore this possibility further in a complementary analysis, we also designed a model in which a single modulated regressor was added to the design matrix, representing all trials collapsed across conditions. To implement this, we constructed design matrices with condition-specific constant regressors and one condition-unspecific parametric modulator representing all trials within a run. All other specification and estimation procedures were identical to those used in the model described above. This estimation was performed for all kinematic features and experiments.

#### 2.4.3. Multivariate pattern analysis (MVPA)

We used MVPA to find the brain regions that code kinematic features of passive finger movement, controlled for the influence of the stimulus duration. For the first-level multivariate analysis, we used the activation beta maps for each unique combination of kinematic features within a run obtained for the unsmoothed functional images. This method increases classification performance and power in MVPA and it is a recommended procedure for event-related experiments (Hebart et al., 2015; Mourão-Miranda et al., 2007, 2006; Mumford et al., 2012). Here, we used The Decoding Toolbox (TDT) (Hebart et al., 2015) for MVPA, written for MATLAB and using SPM functions. We used the searchlight approach with an 8-mm-radius sphere. The searchlight approach trains and test a classifier considering all voxels within a roving region-of-interest (ROI) centered on each voxel and assigning the prediction value to that voxel, until all the voxels in the brain are covered. We used a leave-one-run-out cross-validation for the classification analyses, in which we trained a model with three runs (a cross-validation fold) and validated its prediction performance of the condition level in the remaining run. The classification is performed iteratively for all runs and the average performance is assigned to the central voxel of the searchlight ROI, therefore creating a map of cross-validation accuracies based on the local information around each voxel (Kriegeskorte et al., 2006). To avoid overfitting, the classification was performed through a support-vector machine, i.e., by finding an optimal hyperplane constituted by voxels (features) to define the decision boundary between conditions (classes) (Cortes and Vapnik, 1995). All data was scaled in advance between 0 and 1, which is the recommendation for ‘LIBSVM’ (Chang and Lin, 2007). For each subject, experiment, and kinematic feature, we obtained whole-brain maps of accuracy minus chance (in which the chance was 33.33% (3-class classification) for amplitude and velocity in Exp 1 and 50% (2-class classification) for direction in Exp 1 and for all the conditions in Exp 2). Next, we spatially smoothed the resulting individual maps using a 6-mm isometric Gaussian kernel.

Next, the individual accuracy maps were included in second-level random-effects analyses to compute one-sample t-tests. We thus obtained group-level t-maps in which positive values represent accuracies above chance level for each kinematic feature and experiment. To identify the significant clusters, we used a voxel-wise statistical threshold of p < 0.05, corrected for multiple comparisons using the family-wise error (FWE) method. For Exp 2 and the amplitude condition, we used a voxel-wise statistical threshold of p < 5×10^-5^, corrected for multiple comparisons using the FWE method, because the statistical threshold used for other maps resulted in too many significant voxels for this analysis. An explicit mask was included, considering the whole cerebrum and the superior part of the cerebellum. Using bspmview (github.com/spunt/bspmview), we generated inflated surface maps and reported peak coordinates (multiple peaks were reported for the same clusters if the separation between them was greater than 20 mm), automatically labeled according to the AAL3 atlas.

#### 2.4.4. Univariate analysis

In parallel, we used univariate analysis to find the brain regions showing differential activity across kinematic features of passive finger movement, controlling for stimulus duration, for both experiments. For this first-level analysis, we estimated the activation beta maps for each unique combination of kinematic features within each run using the smoothed functional images. We then created individual linear combination contrast maps for each subject, replicated and scaled across runs, based on the Kronecker product (using the function ‘spm_make_contrasts’) (Henson, 2015) for a three-way repeated-measures Analysis of Variance (ANOVA).

Next, these individual linear combination contrast maps were included in second-level random-effects analyses to compute group-level maps of the main effects of amplitude, velocity, and direction for both experiments separately. This was done using one-way ANOVAs with unequal variance, constituting an appropriate approach for within-subject designs (Henson, 2015). We obtained group-level F-maps, in which values represent activation estimates greater than the grand mean for each kinematic feature and experiment. To identify significant clusters, we used the same voxel-wise statistical thresholds for each condition and experiment and the same explicit mask as in MVPA (section 2.4.3). Inflated surface maps, peak coordinates, and region labels were obtained as in the MVPA.

#### 2.4.5. ROI analyses

To further understand the whole-brain results, we performed ROI analyses centered on the peak coordinates of group accuracy maps for MVPA and group F-value maps for univariate analysis. 5-mm-radius spherical ROIs were created with the MarsBaR toolbox (marsbar.sourceforge.net/; (Brett et al., 2002)). We considered a maximum of six peaks, sorted by the highest accuracy values and labeled with a valid region according to the Anatomy Toolbox. In addition, we performed ROI analyses in regions expected to be involved in passive finger movement according to previous literature. Therefore, we defined six ROIs centered on the regions reported by Galazzo and colleagues (Table 2 from (Boscolo Galazzo et al., 2014)), i.e., the contralateral S1, the SMA, the bilateral S2, the contralateral thalamus, and the ipsilateral cerebellum. The Talairach coordinates provided in the study were converted to MNI space using the function ‘tal2icbm_spm’ (www.brainmap.org/icbm2tal/). The center coordinates are described in Table 1.

**Table 1.** Center coordinates for ROI analyses from regions related to passive finger movement, converted from (Boscolo Galazzo et al., 2014).

| Region Label | MNI Coordinates |  |  |
| --- | --- | --- | --- |
|  | x | y | z |
| L S1 | -41 | -28 | 61 |
| SMA | -8 | -14 | 62 |
| L S2 | -54 | -25 | 20 |
| L S2 | 54 | -27 | 27 |
| L Thalamus | -17 | -20 | 8 |
| R Cerebellum | 25 | -49 | -26 |

**Table 2.** Whole-brain MVPA and univariate results for the amplitude condition and Exp 1. Only clusters containing 10 or more voxels are reported. The statistics column corresponds to accuracy minus chance for MVPA results and to F-value to univariate analysis results. Brain regions were labeled through the Anatomy Toolbox and after visual inspection (between parentheses).

| Region Label | Extent (voxels) | Statistics | MNI Coordinates |  |  |
| --- | --- | --- | --- | --- | --- |
|  |  |  | x | y | z |
| MVPA results |  |  |  |  |  |
| L Postcentral Gyrus (L S1) | 777 | 14.47 | -39 | -34 | 62 |
| L Supramarginal Gyrus |  | 10.04 | -60 | -40 | 41 |
| L Superior Temporal Gyrus | 112 | 9.54 | -57 | -31 | 17 |
| R Superior Frontal Gyrus | 22 | 7.68 | 33 | -16 | 74 |
| R Superior Temporal Gyrus (R M1) | 11 | 7.47 | 63 | -25 | 14 |
| Univariate analysis results |  |  |  |  |  |
| Left precentral gyrus (L M1) | 165 | 48.31 | -40 | -18 | 60 |
| Left midcingulate cortex (L MCC) | 17 | 44.31 | -10 | -30 | 48 |
| Left postcentral gyrus (L S1) | 27 | 42.42 | -36 | -44 | 66 |
| Left postcentral gyrus (L S1) | 20 | 38.17 | -42 | -34 | 66 |

For each ROI defined from whole-brain peaks or literature coordinates, we performed MVPA in the voxels within the ROI to determine whether the region could correctly significantly discriminate at least one of the kinematic feature levels. We trained the classifiers with the levels of each kinematic feature of interest. A leave-one-run-out cross-validation was conducted to compute accuracy performance. We generated subject-specific confusion matrices of accuracy performance for each ROI, experiment, and kinematic feature. The average performance for each true/predicted value in the confusion matrix was considered different from chance according to a bootstrapping approach of 1×10^5^ permutations of subjects. The significance level was set to 0.05 and corrected for multiple comparisons (i.e., the square of the number of condition levels) according to the Bonferroni method. Accuracy performance was considered higher (or lower) than chance if the value was below (above) the confidence interval.

In addition, for each ROI defined from whole-brain peaks or literature coordinates, we performed post-hoc univariate analyses within each ROI to determine whether the region showed significant pairwise differential activation estimates. First, we extracted the average beta of the activation estimates from each unique combination of kinematic features within each ROI. Next, we performed one-way ANOVAs using RStudio (www.rstudio.com/) to compare activation estimates across condition levels for each ROI, experiment, and kinematic feature. For this analysis, we used the function ‘emmeans_test’ (library ‘rstatix’), with a significance level of 0.05 and correction for multiple comparisons according to the Sidak method.

The complementary analysis (using a single modulated regressor) was performed only for the univariate ROI analysis, using the ROIs defined from the literature coordinates, because we wanted to determine how the general pattern of results in key regions would be altered by a model controlling for the interaction stimulus duration by condition. This complementary analysis was conducted for all kinematic features and experiments. We extracted the average beta of activation estimates from the conditions of interest (constant regressors) within each ROI for each unique combination of kinematic features. All other procedures for this analysis were identical to the main analysis described above.

#### 2.4.6. Evaluation of MVPA stability across subjects

We additionally analyzed the stability of the MVPA models by classifying state levels across subjects. The stability analysis estimates how generalizable the MVPA results are across subjects. This analysis was done for all ROIs defined from the literature coordinates (section 2.4.5) and for kinematic features and experiments separately. We used the activation estimates for each unique combination of kinematic features determined from the unsmoothed functional images, controlling for the influence of stimulus duration. For each condition level, we trained the model using the condition level estimates for all runs from all subjects minus one and tested it on the remaining subject (leave-one-subject-out cross-validation). No scaling method was applied. We obtained the confusion matrices of the mean accuracies across subjects for each kinematic feature and experiment. Since only one confusion matrix was obtained for the group for this analysis, no statistical testing was applied. The MVPA ROI analysis was considered stable across subjects if the main diagonal was clearly higher than the other values in the resulting confusion matrix.

## 3. Results

### 3.1. Amplitude

#### 3.1.1. Experiment 1

For Exp 1 (which included three amplitude levels), whole-brain MVPA revealed a brain network encoding amplitude levels of passive finger movement, which included contralateral S1 and SMG, ipsilateral M1, and bilateral STG (Fig. 2A and Table 2). Whole-brain univariate analysis, for the same statistical threshold, showed differential activation across amplitude conditions in the contralateral M1 and S1 and in the MCC (Fig. 2B and Table 2). The activation estimates in these regions increased with amplitude. Overall, whole-brain MVPA was more spatially sensitive than univariate analysis for amplitude of passive movement. MVPA ROI analysis for Exp 1 further showed that, in addition to contralateral S1, the contralateral and ipsilateral S2 also encode amplitude levels (Fig. 2C). Univariate ROI analysis showed that, in addition to contralateral S1, the thalamus and cerebellum showed differential activation across amplitude levels (Fig. 2D), but not bilateral S2. There was no acceptable stability for amplitude in Exp 1 (Fig. S1A, top row). Axial slices for whole-brain maps for amplitude in Exp 1 obtained with MVPA and univariate analysis are shown in Fig. S2.

**Figure 2.**
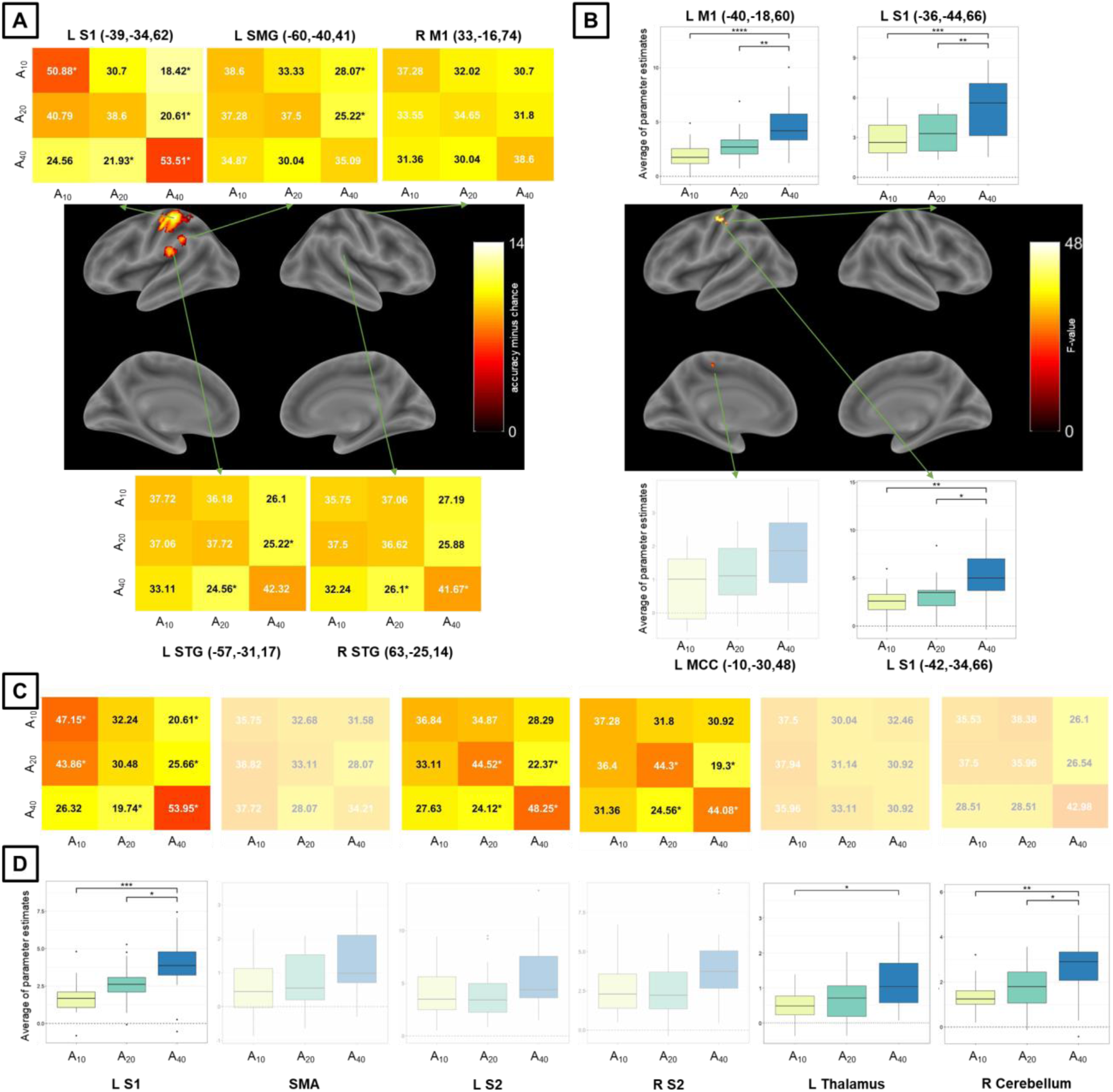
MVPA and univariate analysis results for the amplitude condition and Exp 1. Whole-brain results for (A) MVPA (accuracy significantly above chance) and (B) univariate analysis (significant differential activation), both using a voxel-level inclusion threshold of p < 0.05, FWE (family-wise error)-corrected for multiple comparisons. ROI analysis results (C) for MVPA and (D) univariate analysis. Non-transparent plots in (A) and (C) represent accuracies significantly different from chance. Non-transparent plots in (B) and (D) represent significant main effects of amplitude in activation and post-hoc significant pairwise differences. Asterisks in MVPA results represent accuracies significantly different from chance (33.33%), corrected for multiple comparisons using the Bonferroni method and bootstrapping (10^5^ permutations) across subjects. Asterisks in univariate analysis results represent significant differences across activation estimates, corrected for multiple comparisons (Sidak method – ****: p < 0.0001, ***: p < 0.001, ** p < 0.01, * p < 0.05). For MVPA confusion matrices, numbers in black (white) are higher (lower) than chance. S1/S2 = primary/secondary somatosensory cortex, SMA = supplementary motor area, SMG = supramarginal gyrus, M1 = primary motor cortex, STG = superior temporal gyrus, MCC = midcingulate cortex, L/R = left/right.

#### 3.1.2. Experiment 2

For Exp 2 (which included two amplitude levels), whole-brain MVPA revealed a large brain network encoding amplitude levels of passive finger movement, which included the bilateral sensorimotor network (including S1 and S2, M1, and SMA) (Fig. 3A and Table 3). Whole-brain univariate analysis, for the same statistical threshold, showed differential activation across amplitude conditions in the contralateral S1 and S2, and the ipsilateral STG (Fig. 3B and Table 3). The activation estimates in these regions increased with amplitude. MVPA for Exp 2 showed high sensitivity, despite the strict statistical threshold compared to the analysis for amplitude in Exp 1. MVPA ROI analysis for Exp2 further showed that all analyzed regions encode amplitude levels (Fig. 3C). Univariate ROI analysis also showed that all analyzed regions exhibited differential activation across amplitude levels (Fig. 3D). Stability was acceptable for amplitude in Exp 2 (Fig. S1A, bottom row). Axial slices for whole-brain maps for amplitude in Exp 2 obtained with MVPA and univariate analysis are shown in Fig. S3.

**Figure 3.**
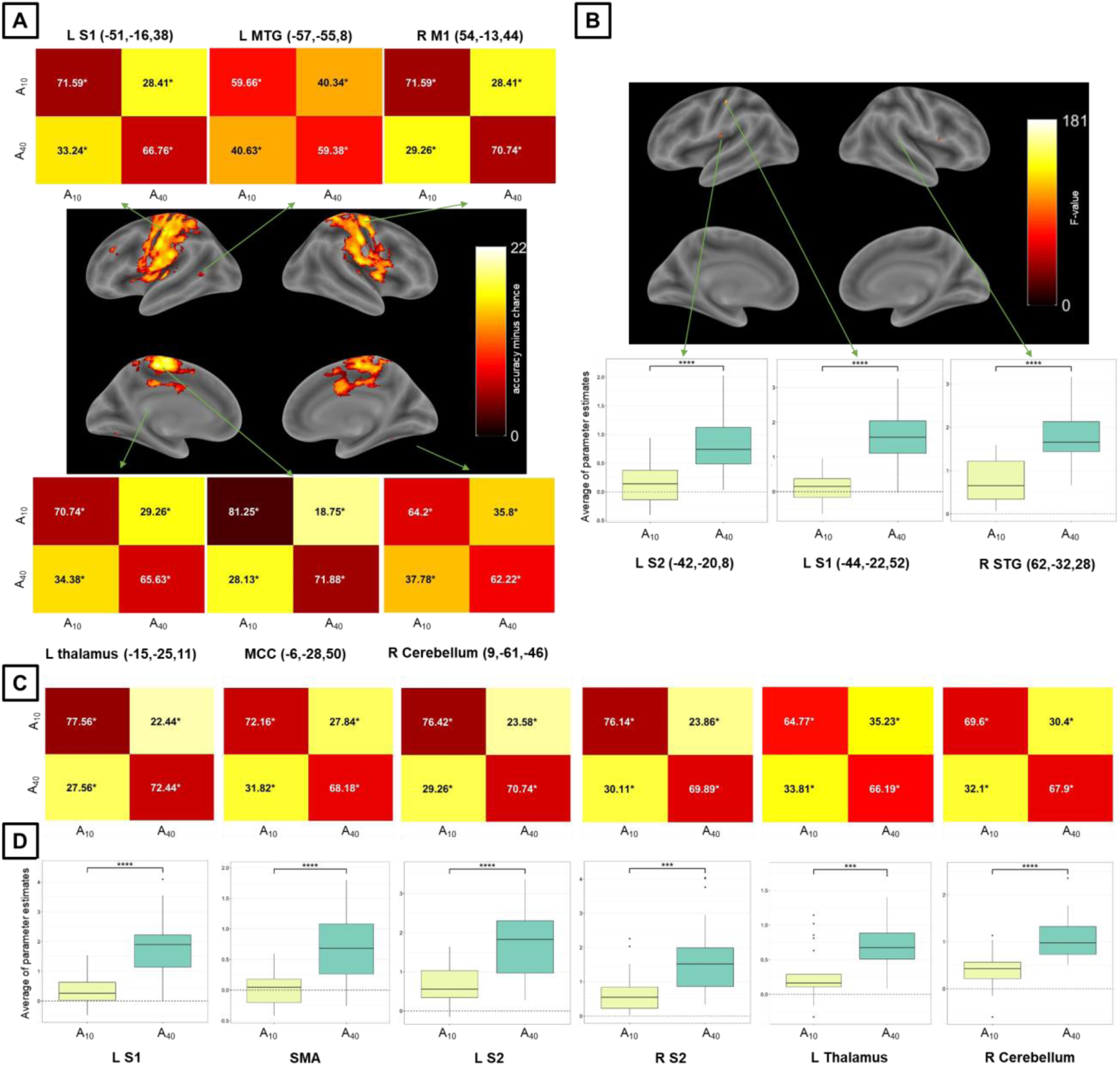
MVPA and univariate analysis results for the amplitude condition and Exp 2. (A) Whole-brain results for (A) MVPA (accuracy significantly above chance) and (B) univariate analysis (significant differential activation), both using a voxel-level inclusion threshold of p < 5×10^-5^, FWE (family-wise error)-corrected for multiple comparisons. ROI analysis results (C) for MVPA and (D) univariate analysis. Plots in (A) and (C) represent accuracies significantly different from chance. Plots in (B) and (D) represent significant main effects of amplitude in activation and post-hoc significant pairwise differences. Asterisks in MVPA results represent accuracies significantly different from chance (50%), corrected for multiple comparisons using the Bonferroni method and bootstrapping (10^5^ permutations) across subjects. Asterisks in univariate analysis results represent significant differences across activation estimates, corrected for multiple comparisons (Sidak method – ****: p < 0.0001, ***: p < 0.001). For MVPA confusion matrices, numbers in black (white) are higher (lower) than chance. S1/S2 = primary/secondary somatosensory cortex, MTG = middle temporal gyrus, M1 primary motor cortex, MCC = midcingulate cortex, STG = superior temporal gyrus, SMA = supplementary motor area, L/R = left/right.

**Table 3.** Whole-brain MVPA and univariate results for the amplitude condition and Exp 2. Only clusters containing 10 or more voxels are reported. The statistics column corresponds to accuracy minus chance for MVPA results and to F-value to univariate analysis results. Brain regions were labeled through the Anatomy Toolbox and after visual inspection (between parentheses).

| Region Label | Extent<br>(voxels) | Statistics | MNI Coordinates |  |  |
| --- | --- | --- | --- | --- | --- |
|  |  |  | x | y | z |
| MVPA results |  |  |  |  |  |
| L MCC | 6400 | 22.78 | -6 | -28 | 50 |
| L Postcentral Gyrus (L S1) |  | 19.33 | -51 | -16 | 38 |
| R Postcentral Gyrus (R S1) |  | 19.25 | 54 | -13 | 44 |
| L Thalamus | 33 | 16.23 | -15 | -25 | 11 |
| R Cerebellum (IX) | 86 | 15.52 | 9 | -61 | -46 |
| R Cerebellum (VI) | 68 | 13.81 | 30 | -52 | -25 |
| L Middle Temporal Gyrus | 20 | 13.81 | -57 | -55 | 8 |
| L IFG (p. Triangularis) | 17 | 13.30 | -39 | 29 | 29 |
| L Fusiform Gyrus | 11 | 12.57 | -27 | -64 | -1 |
| Univariate analysis results |  |  |  |  |  |
| L Insula Lobe (L S2) | 50 | 174.68 | -42 | -20 | 8 |
| L Postcentral Gyrus (L S1) | 25 | 181.72 | -44 | -22 | 52 |
| R Supramarginal Gyrus (R SMG) | 10 | 155.71 | 62 | -32 | 28 |

### 3.2. Velocity

#### 3.2.1. Experiment 1

For Exp 1 (which included three velocity levels), neither the whole-brain MVPA nor the whole-brain univariate analysis led to surviving clusters related to velocity of passive finger movement. Neither the MVPA nor the univariate ROI analysis showed significant results for velocity in Exp 1 (Fig. S4). There was no acceptable stability for velocity in Exp 1 (Fig. S1B, top row).

#### 3.2.2. Experiment 2

For Exp 2 (which included two velocity levels), whole-brain MVPA resulted in no surviving clusters for velocity of passive finger movement. Whole-brain univariate analysis showed, for the same statistical threshold, showed greater activity in the contralateral S1 for a lower compared to a higher velocity, as well as lower activity in the precuneus for a higher compared to a lower velocity (Fig. 4A and Table 4). MVPA and univariate ROI analysis for Exp 2 showed that the SMA was the only region, among the ones analyzed, to both encode (Fig. 4B) and exhibit differential activation (Fig. 4C) for velocity. There was no acceptable stability for velocity in Exp 2 (Fig. S1B, bottom row). Axial slices for whole-brain maps for velocity in Exp 2 obtained with univariate analysis are shown in Fig. S5.

**Figure 4.**
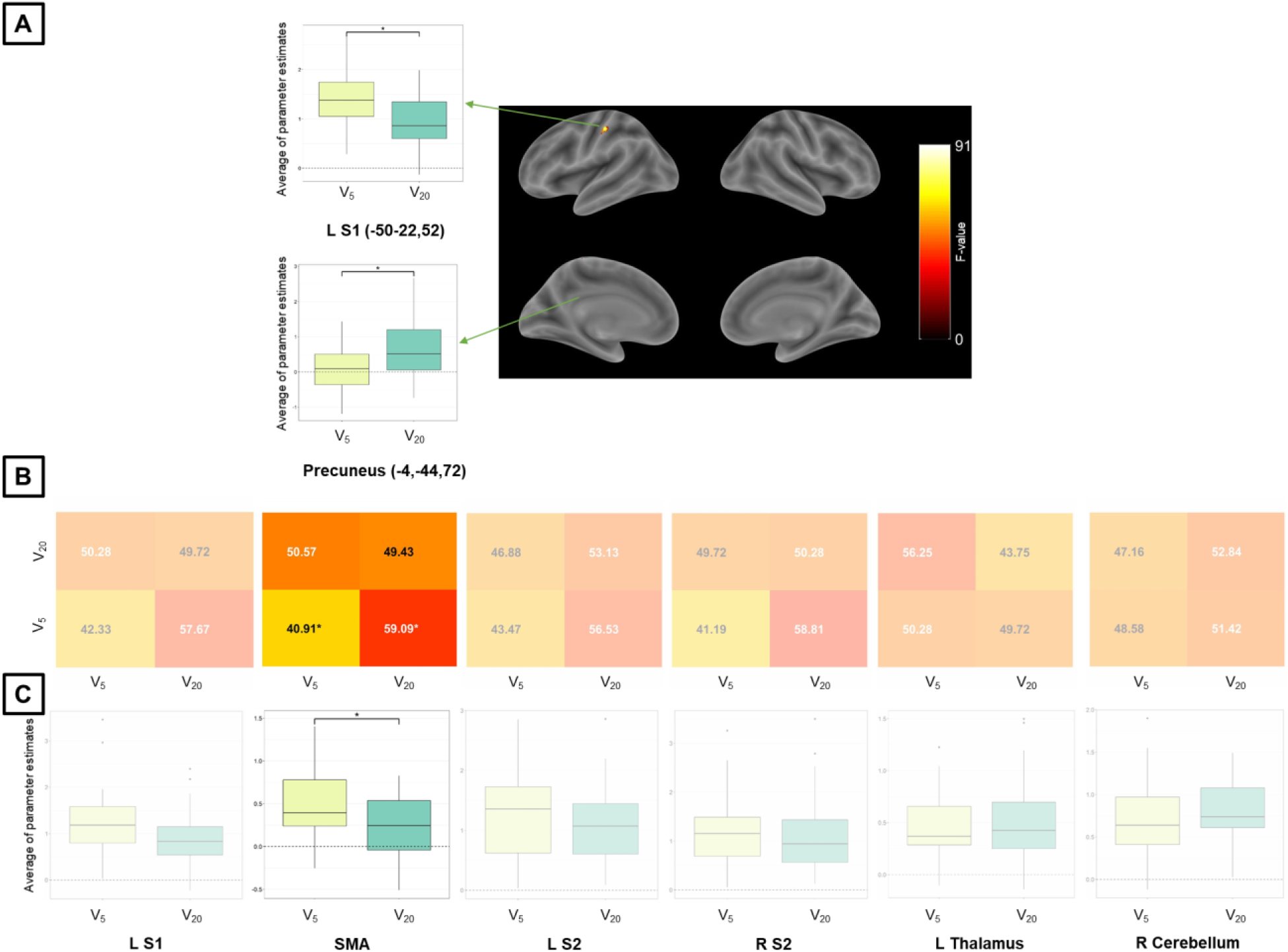
MVPA and univariate analysis results for the velocity condition and Exp 2. (A) Whole-brain results for univariate analysis (significant differential activation), using a voxel-level inclusion threshold of p < 0.05, FWE (family-wise error)-corrected for multiple comparisons. ROI analysis results (B) for MVPA and (C) univariate analysis. The non-transparent plot in (B) represents accuracies significantly different from chance. The non-transparent plot in (C) represents significant main effects of amplitude in activation and a post-hoc significant pairwise difference. The asterisks in the MVPA result represents accuracy significantly different from chance (50%), corrected for multiple comparisons using the Bonferroni method and bootstrapping (10^5^ permutations) across subjects. Asterisks in univariate analysis results represent significant differences, corrected for multiple comparisons (Sidak method – *: p < 0.05). For MVPA confusion matrices, numbers in black (white) are higher (lower) than chance. S1/S2 = primary/secondary somatosensory cortex, SMA = supplementary motor area, L/R = left/right.

**Table 4.** Whole-brain univariate results for the velocity condition and Exp 2. Only clusters containing 10 or more voxels are reported. Brain regions were labeled through the Anatomy Toolbox and after visual inspection (between parentheses).

| Region Label | Extent (voxels) | F-value | MNI Coordinates |  |  |
| --- | --- | --- | --- | --- | --- |
|  |  |  | x | y | z |
| L Postcentral Gyrus (L S1) | 79 | 91.03 | -50 | -22 | 52 |
| L Precuneus | 24 | 64.64 | -4 | -44 | 72 |

### 3.3. Direction

#### 3.3.1. Experiment 1

For Exp 1, whole-brain MVPA revealed a brain network encoding direction levels of passive finger movement, which included contralateral S1, M1, and STG, ipsilateral SMG, and MCC and precuneus (Fig. 5A and Table 5). Whole-brain univariate analysis, for the same statistical threshold, showed higher activation for extension than for flexion in the SMA (Fig. 5B and Table 5). Whole-brain MVPA was more spatially sensitive than univariate analysis for direction of passive movement. MVPA ROI analysis for Exp 1 further showed that, in addition to contralateral S1, the ipsilateral S2 and thalamus also encode direction levels (Fig. 5C). Univariate ROI analysis showed the same result observed for whole-brain analysis (Fig. 5D). There was no acceptable stability for direction in Exp 1 (Fig. S1C, top row). Axial slices for whole-brain maps for direction in Exp 1 obtained with MVPA and univariate analysis are shown in Fig. S6.

**Figure 5.**
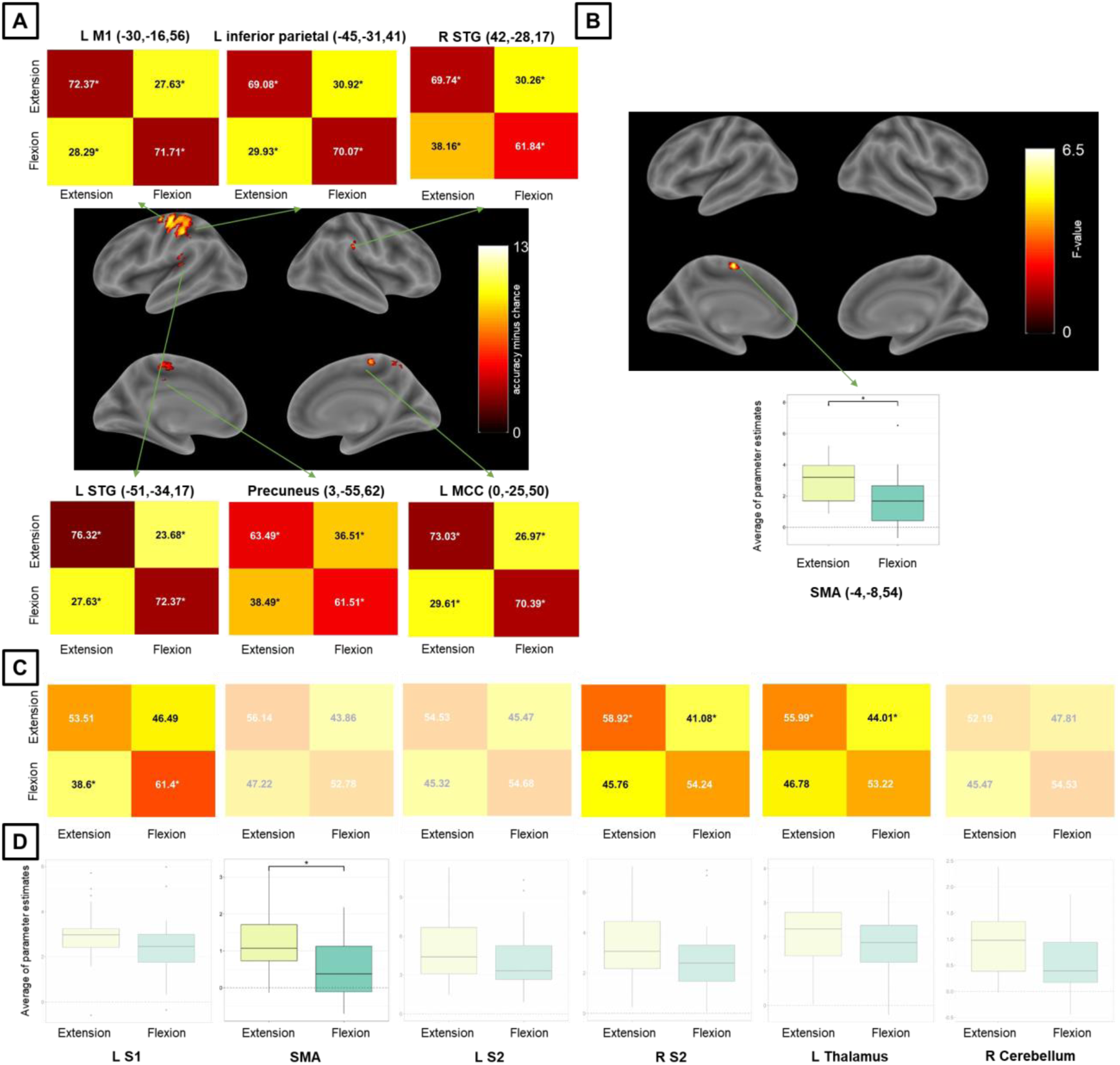
MVPA and univariate analysis results for the direction condition and Exp 1. Whole-brain results for (A) MVPA (accuracy significantly above chance) and (B) univariate analysis (significant differential activation), both using a voxel-level inclusion threshold of p < 0.05, FWE (family-wise error)-corrected for multiple comparisons. ROI analysis results (C) for MVPA and (D) univariate analysis. Non-transparent plots in (A) and (C) represent accuracies significantly different from chance. Non-transparent plots in (B) and (D) represent significant main effects of amplitude in activation and post-hoc significant pairwise differences. Asterisks in MVPA results represent accuracies significantly different from chance (50%), corrected for multiple comparisons using the Bonferroni method and bootstrapping (10^5^ permutations) across subjects. Asterisks in univariate analysis results represent significant differences across activation estimates and are corrected for multiple comparisons (Sidak method – *: p < 0.05). For MVPA confusion matrices, numbers in black (white) are higher (lower) than chance. S1/S2 = primary/secondary somatosensory cortex, MTG = middle temporal gyrus, M1 = primary motor cortex, MCC = midcingulate cortex, SMA = supplementary motor area, L/R = left/right.

**Table 5.**
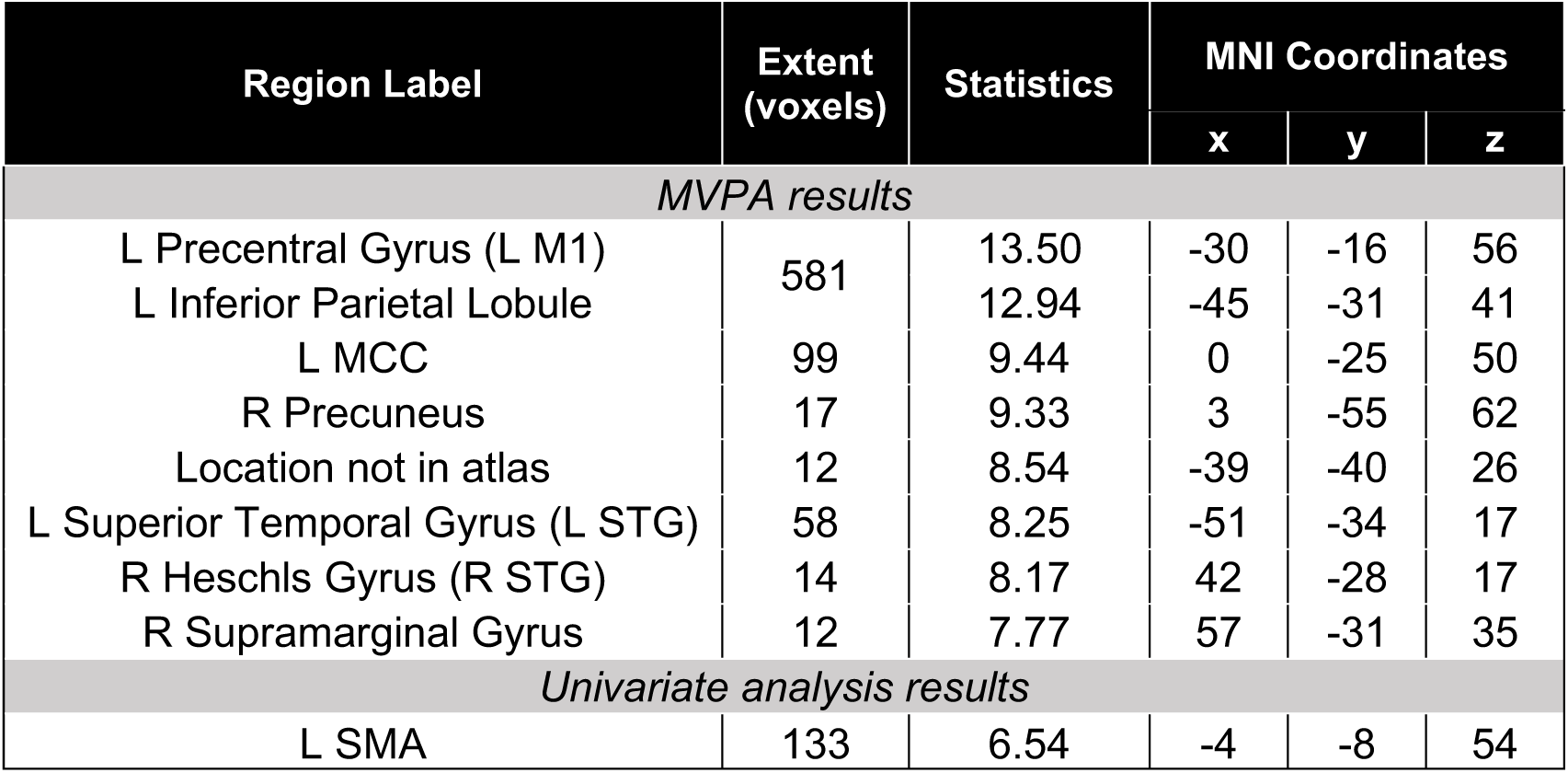
Whole-brain MVPA and univariate results for the direction condition and Exp 1. Only clusters containing 10 or more voxels are reported. The statistics column corresponds to accuracy minus chance for MVPA results and to F-value to univariate analysis results. Brain regions were labeled through the Anatomy Toolbox and after visual inspection (between parentheses).

| Region Label | Extent (voxels) | Statistics | MNI Coordinates |  |  |
| --- | --- | --- | --- | --- | --- |
|  |  |  | x | y | z |
| MVPA results |  |  |  |  |  |
| L Precentral Gyrus (L M1) | 581 | 13.50 | -30 | -16 | 56 |
| L Inferior Parietal Lobule |  | 12.94 | -45 | -31 | 41 |
| L MCC | 99 | 9.44 | 0 | -25 | 50 |
| R Precuneus | 17 | 9.33 | 3 | -55 | 62 |
| Location not in atlas | 12 | 8.54 | -39 | -40 | 26 |
| L Superior Temporal Gyrus (L STG) | 58 | 8.25 | -51 | -34 | 17 |
| R Heschls Gyrus (R STG) | 14 | 8.17 | 42 | -28 | 17 |
| R Supramarginal Gyrus | 12 | 7.77 | 57 | -31 | 35 |
| Univariate analysis results |  |  |  |  |  |
| L SMA | 133 | 6.54 | -4 | -8 | 54 |

#### 3.3.2. Experiment 2

For Exp 2, neither the whole-brain MVPA nor the whole-brain univariate analysis led to surviving clusters for the direction of passive finger movement. MVPA ROI analysis for Exp 2 showed that the contralateral S1 was the only region to encode direction levels (Fig. 6A). There were no significant results for univariate ROI analysis (Fig. 6B) and no acceptable stability for direction in Exp 2 (Fig. S1C, bottom row).

**Figure 6.**
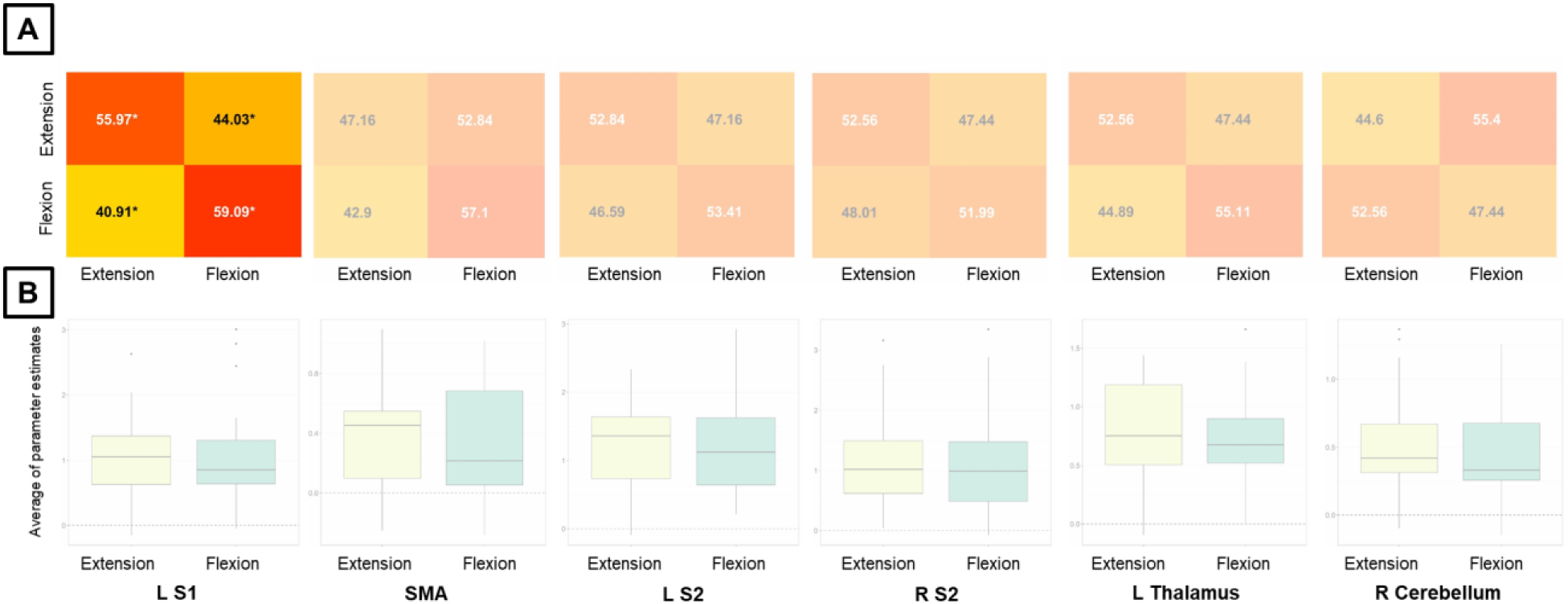
MVPA and univariate analysis results for the direction condition and Exp 2. ROI analysis results (A) for MVPA and (B) univariate analysis. The non-transparent plot in (A) represents accuracies significantly different from chance. Asterisks in MVPA results represent accuracies significantly different from chance (50%), corrected for multiple comparisons using the Bonferroni method and bootstrapping (10^5^ permutations) across subjects. For MVPA confusion matrices, numbers in black (white) are higher (lower) than chance. S1/S2 = primary/secondary somatosensory cortex, SMA = supplementary motor area, L/R = left/right.

## 4. Discussion

We provide evidence for the brain regions that respond to different kinematic elements of passive finger movement using fMRI data from neurotypical subjects. In two experiments, we investigated how kinesthesia is encoded in the brain by using an MRI-compatible robot to provide subjects with systematic finger stimulation. Our analyses consisted of univariate and multivariate approaches, performed separately for each kinematic feature and using a model that accounts for the influence of stimulus duration. This model mitigates the false positive rates regardless of whether neural activity scales with stimulus duration. Previous research on muscle spindles and skin receptors in kinesthesia has suggested that proprioception can be dissociated into position, velocity, and direction senses (Proske and Gandevia, 2009). Therefore, such evidence has led to the hypothesis that kinematic features might also be spatially dissociated in the brain. We confirmed previous findings that a number of somatosensory and motor regions are related to the amplitude and direction of passive finger movement. In contrast, we observed fewer brain regions associated with the perception of velocity. Thus, we provide evidence that brain responses to passive finger movements are primarily reliant on amplitude, i.e., the change in finger positioning relative to a starting point. Other kinematic features are also represented in the brain, but to a lesser extent. As we will discuss in detail in the following sections, the finding that brain responses are primarily sensitive to changes in position, rather than to the rate or direction of these spatial changes, suggests that passive movement relies mainly on the measurement of finger position through muscle spindles and on the constant updating of expected position.

### 4.1. Interpretation of univariate and multivariate results in fMRI

Univariate and multivariate approaches based on the high spatial resolution of fMRI can be used to study brain regions involved in psychological or physical aspects. A univariate approach estimates the extent to which the BOLD time-course of a single voxel fits a predefined model for a given condition of interest. Thus, higher activation is interpreted as a higher level of processing or resource utilization for one condition compared to another in a brain region. This approach, however, can be insensitive to processing patterns of activity that follow a set of distributed voxels (Davis et al., 2014). A multivariate approach to fMRI brain mapping considers patterns of activity distributed across voxels, rather than separately within individual voxels, and trains a model using a machine-learning algorithm to classify patterns into psychological or physical conditions. The model’s successful classification performance then reflects that the brain region is associated with the condition in question. MVPA is more sensitive in mapping functional correlates of brain regions compared to univariate approaches (Coutanche, 2013; Davis and Poldrack, 2013; Kriegeskorte et al., 2006; Norman et al., 2006), and is therefore a powerful approach for investigating how the brain encodes behaviors of interest.

It is hence crucial to properly interpret the different outcomes of univariate and multivariate approaches to fMRI. It is possible that a univariate approach shows no activation for a region while that region contains multivariate information (Harrison and Tong, 2009), meaning that the condition of interest is represented across distributed voxels rather than spatially localized. On the other hand, the presence of a univariate activity difference means that the condition of interest can be differentiated by activity patterns with a lower spatial variance (Coutanche, 2013). It has been suggested that multivoxel patterns reflect representational content within a region (Mur et al., 2009; Riggall and Postle, 2012), but MVPA does not necessarily reveal details about the dimensionality of a representation (Davis et al., 2014). Essentially, it is thought that the increased sensitivity of MVPA derives from greater spatial heterogeneity that is undetectable by a univariate approach, and from neglecting subject-level variability accounted for in univariate analyses (Coutanche, 2013; Davis et al., 2014). Here, we applied both approaches to understand and complement the neural basis of kinematic passive movement features.

### 4.2. Amplitude of passive finger movement

We investigated brain regions associated with amplitude of passive finger movement. We confirmed previous findings of a network comprised of sensorimotor and subcortical regions associated with amplitude of passive finger movement and other body parts in humans (Boscolo Galazzo et al., 2014; Francis et al., 2009; Jaeger et al., 2014; Mima et al., 1999; Szameitat et al., 2012; Weiller et al., 1996). Overall, univariate analysis confirmed that activity in these regions increases with amplitude, concordant with studies on active movement (Shirinbayan et al., 2019) and afferent firing rates (Edin, 1992; Knibestöl, 1975). Compared to other kinematic features, we associated amplitude of passive movement with strong univariate differences across conditions and large discriminative areas in multivariate analysis. Limb position is first detected by muscle spindles (Goodwin et al., 1972) and skin receptors (Edin, 1992) and this information is then carried to the brain. Static discharges that proportionally change according to stretch of muscle spindles (Roll and Vedel, 1982). Stronger brain activity in response to amplitude than to velocity seems congruent with evidence that information about the sense of position comes from muscle spindles. Although the primary endings of muscle spindles respond to both length and velocity changes, the secondary endings are only sensitive to length change (Matthews, 1974; Proske and Gandevia, 2009). As an alternative interpretation, during passive movement, the final position is unknown and the expectation of position is continuously updated by proprioception until the movement ends. Greater proprioceptive updating of positional expectation would therefore be required for estimating larger displacements during passive movement. Our results suggest that brain activation in the somatosensory cortex scales with proprioceptive inputs from passive finger movement, leading to an association between somatosensory activity and amplitude of passive movement, independent of stimulus duration.

We observed an association between contralateral S1 activity and amplitude of passive finger movement in Exps 1 and 2 (Figs. 2 and 3), which is consistent with previous studies on passive and active movement (Boscolo Galazzo et al., 2014; Guzzetta et al., 2007; Lolli et al., 2021, 2019; Shirinbayan et al., 2019; Weiller et al., 1996; Zhavoronkova et al., 2017). S1 activity was also observed in early studies of cortical recordings in nonhuman primates during passive finger movement (Gardner and Costanzo, 1981). In addition, we observed ipsilateral S1 activity associated with amplitude, but only for MVPA in Exp 2 (Fig. 3A). The S1 activity increasing with amplitude is consistent with previous studies on active and passive finger movement (Nurmi et al., 2018; Shirinbayan et al., 2019). The proprioceptive and somatosensory information that S1 receives during movement (Mountcastle et al., 1992; Yousry et al., 1997) may provide finger localization during passive movement. We also observed bilateral S2 activity associated with amplitude, for MVPA in Exp 1 (Fig. 2A,C) and for both MVPA and univariate analysis in Exp 2 (Fig. 3). S2 has been associated with passive finger movement (Dueñas et al., 2018; Mima et al., 1999), but less sensitive than S1 (Backes et al., 2000; Nelson et al., 2004; Nurmi et al., 2018). While S1 receives rather simple peripheral sensory information, S2 may be related to higher aspects of sensory processing and retention of relevant afferent aspects in working memory (Lemus et al., 2010). For Exp 1, we observed results in S2 only for MVPA (Fig. 2C), which may reflect that the amplitude-related S2 activity is more spatially distributed compared with S1.

We observed that the SMA, the contralateral SMG, the posterior part of the STG, and the contralateral M1 were also involved in the amplitude of passive finger movement (Figs. 2 and 3). The SMA is involved in movement preparation, planning, and control (Tanji, 1994) and has been reported to be associated with passive movement (Alary et al., 1998; Carel et al., 2000; Guzzetta et al., 2007; Nurmi et al., 2018; Weiller et al., 1996). As reported in other studies on active and passive movement, we confirmed that SMA activation was more dominant in the contralateral hemisphere (Pool et al., 2013; Zhavoronkova et al., 2017). It has been suggested that the SMA receives information from the somatosensory system (Cadoret and Smith, 1997) and its activation (as well as the contralateral S1) is lower during passive compared to active movement (Lolli et al., 2021; Mima et al., 1999; Zhavoronkova et al., 2017). Thus, the SMA would be more sensitive to afferent information during active than passive movement and would differentiate between intentional and involuntary movement (Mima et al., 1999). In a different vein, the SMG is part of the somatosensory cortex, processes proprioceptive information (Ben-Shabat et al., 2015), and is involved in spatial perception (Bjoertomt et al., 2009). In addition, M1 is known to be primarily involved in motor execution of body parts. Consistent with our results, single-cell recordings from non-human primates (Schwartz, 1992), and estimation of BOLD activity (Shirinbayan et al., 2019) showed that M1 is associated with movement amplitude, but not velocity (Section 3.2). Interestingly, while M1 remains silent during tactile stimulation, it is active during both active and passive movement (Goldring and Ratcheson, 1972). Therefore, M1 is thought to be involved in proprioceptive feedback independent of skin stimulation (Jansma et al., 1998).

Besides the cortical regions, we also observed that the thalamus and the ipsilateral cerebellum were involved in the amplitude of passive finger movement (Fig. 2 and 3). In Exp 1, the thalamus and ipsilateral cerebellum was associated with amplitude only for the univariate analysis (Fig. 2D). That the thalamus and cerebellum are primarily involved in differential activity, rather than fine coding, may be due to their less specialized involvement, compared to the higher aspects of sensory processing in S2. The thalamus has been associated with active and passive movement, more prominently in the contralateral hemisphere (Boscolo Galazzo et al., 2014; Shirinbayan et al., 2019), and sends direct projections to the somatosensory cortex (Viaene et al., 2011). Ipsilateral cerebellar responses to passive movement have also been reported (Boscolo Galazzo et al., 2014; Zhavoronkova et al., 2017). Passive movement features may be processed in these regions, likely due to afferent nerves in the brainstem reaching both the cerebellum and the cortex (the latter, crossing the thalamus) (Landgren and Silfvenius, 1971; Proske and Gandevia, 2009).

### 4.3. Velocity of passive finger movement

We also examined brain regions associated with the velocity of passive finger movement. We observed that sensorimotor network regions (contralateral S1 and SMA) showed a negative relationship between activation and velocity, whereas the precuneus showed a positive relationship with velocity. Compared to amplitude, fewer cortical brain regions were associated with velocity during passive movement (Fig. 4), which is consistent with previous evidence from spike train recordings in animals (Golub et al., 2014) and studies in humans using fMRI (Shirinbayan et al., 2019; Wenzel et al., 2014) during active movement. In addition, whole-brain univariate analysis yielded significant results in sensorimotor regions that, surprisingly, were not found in whole-brain MVPA.

Univariate activity in contralateral S1 and SMA was associated with velocity of passive movement, consistent with previous studies (Boscolo Galazzo et al., 2014; Dueñas et al., 2018; Mima et al., 1999; Szameitat et al., 2012). A study on active movement showed that the contralateral S1 is more specific for amplitude than for velocity, congruent to our hypothesis, whereas the SMA is equally involved in both amplitude and velocity (Shirinbayan et al., 2019). The peak coordinates for the main effect of velocity were more frontal than for amplitude. In contrast to S1, we observed no evidence for S2 involvement with velocity, similar to findings using nerve stimulation (Backes et al., 2000) or peripheral stimulation (Nelson et al., 2004). We found a negative relationship between velocity and activation in the contralateral S1 and SMA. This inverse relationship between activity and velocity, independent of stimulus duration, may be due to increased proprioceptive updating during slower trajectories of passive movement. Therefore, we suggest that proprioceptive afferent information during passive movement may depend on velocity, with slower trajectories carrying more proprioceptive information.

Some studies have reported the relationship between velocity of passive movement and brain activity based on oscillatory frequencies. Here, we argue that this movement pattern is fundamentally different from linear displacement and may not be directly comparable to our results. During passive linear displacement, the fingers start and end at different positions in space, and proprioception plays a more critical role in spatial localization than in the case of oscillatory stimulation. This experimental difference may be the reason why studies on passive and active movement reported that the sensorimotor cortex and cerebellum were strongly related to movement rate (Nurmi et al., 2018; Riecker et al., 2003; Taniwaki et al., 2003; Turner et al., 1998) while our results with linear velocity were less sensitive. Similar to our study, Shirinbayan et al. (2019) found less sensitive results for velocity of linear displacements during active movements compared to amplitude. It is also possible that, during rapid oscillatory movements, the neural mechanisms of proprioception for frequency and amplitude are blended. For instance, an early study showed that muscle vibration, initially perceived as movement during rapid oscillatory vibration, was perceived as a change in position at slower frequencies (McCloskey, 1973), with position and velocity dissociated in muscle spindles. In fact, it has been proposed that a moderate oscillation rate (and therefore dissociation of position and velocity) is optimal to generate the greatest activation in sensorimotor cortex (Bae et al., 2017). Conversely, the univariate approach showed that the precuneus was positively related to the velocity of passive movement. The precuneus has also been implicated in passive movement in previous studies (Jaeger et al., 2014; Sahyoun et al., 2004). The precuneus is part of a network that is activated by higher levels of internally-focused attention (Andrews-Hanna et al., 2014). Therefore, we can only speculate that the faster passive movement may have induced heightened internally-oriented attention and, consequently, elicited precuneus activity.

### 4.4. Direction of passive finger movement

Finally, we also investigated brain regions associated with the direction of passive finger movement, i.e., extension or flexion. We found a set of regions associated with passive movement direction located in the somatosensory cortex, roughly similar to the regions associated with amplitude. We found that contralateral S1 and M1, bilateral STG, ipsilateral S2, SMA, and thalamus were associated with passive movement direction (Figs. 5 and 6). From the univariate analysis, we found that SMA activity was higher during extension than flexion. Results regarding movement direction are less common in the literature compared to other kinematic features, with some studies reporting no findings for direction (Dueñas et al., 2018; Shirinbayan et al., 2019).

The contralateral S1 related to the direction of passive movement was identified only by MVPA and not by the univariate approach. This finding may imply that the spatial representation for direction is diffuse rather than locally concentrated, which differs from our amplitude and velocity results. In fact, Hammer et al. suggested that directional tuning is better represented neurally by large populations of neurons (Hammer et al., 2016); Eisenberg et al. also showed that directional tuning follows a complex spatial organization (Eisenberg et al., 2010), which is likely lost in second-level group analyses. Previous evidence suggests that directional afferents are primarily based on skin receptors (Proske and Gandevia, 2009). Here, we speculate that directional afferent information from skin receptors may have a more diffuse brain representation than amplitude, but further studies are needed to confirm this hypothesis. Clinically, when the proprioception is assessed, patients are typically asked to indicate the direction of a passive movement (usually, up or down) (Lincoln et al., 1998). Our results indicate that the amplitude of passive finger movement is more broadly represented in the brain, suggesting that this kinematic feature could also be advantageously used to design more sensitive clinical assessments (Zbytniewska et al., 2021).

### 4.5. Methodological considerations

Here we would like to address some points related to the modeling strategy used in this study. Of all the combinations of experiments and kinematic features, only the data from Exp 2 for amplitude led to acceptable stability (Fig. S1A, bottom row), i.e., the model trained with these data was reproducible across subjects. We argue that the increased sensitivity combined with the reduced number of levels was important for achieving acceptable stability. Furthermore, Exp 2 yielded results with higher sensitivity to velocity than Exp 1 (Figs. S4 and 4), thus fewer levels may be appropriate to study the brain correlates of velocity. However, in contrast to amplitude and velocity, data from Exp 1 yielded more sensitive results for direction than Exp 2. This discrepancy may be because Exp 2 had fewer trials available to estimate run-specific betas than Exp 1.

Amplitude, velocity, and stimulus duration are inherently related, and varying one feature inevitably implies varying another. In block designs, varying velocity, for example, implies that data will be confounded with frequency or number of repetitions (Shirinbayan et al., 2019). In an event-related design, varying, for example, amplitude implies that velocity or stimulus duration will also vary (Turner et al., 2003). Therefore, it is desirable to control results for stimulus duration when studying brain activation related to physical movement. Time-on-task is positively correlated with BOLD activation (Taylor et al., 2014; Yarkoni et al., 2009) and can confound the results (Mumford et al., 2015). While reaction time in cognitive neuroscience may carry relevant information (Pamplona et al., 2022; Woolgar et al., 2014), stimulus duration in physical stimulation may be of less physiological interest. Stimulus duration can also potentially be a confounder in MVPA, i.e., the classification will also identify brain regions that are associated not only with a condition of interest, but also with stimulus duration (Todd et al., 2013). In fact, MVPA is very sensitive to any differences (Woolgar et al., 2014), and thus addressing confounders is important to obtain reliable results and accurate interpretations.

The first study to analyze the present dataset (Dueñas et al., 2018) estimated activation using variable epochs (Grinband et al., 2008) to account for stimulus duration in the model. This model appropriately scales the estimate when neural activation scales with stimulus duration, but leads to inflated error rates when neural activation does not scale with stimulus duration (Mumford et al., 2024), which may be fundamentally the case when characterizing brain regions that are intrinsically related to sensorimotor components of passive movement rather than simply stimulus duration. In the present study, we constructed the regressors with constant-duration boxcar functions modulated by stimulus duration, which accounts for stimulus duration and mitigates false positives regardless of whether neural activation scales with stimulus duration (Mumford et al., 2024). This methodological difference may explain the differences in univariate results between the seminal study and ours. For instance, our study found a positive relationship between contralateral S1 activation and amplitude, whereas the previous study found a negative relationship. In the current study, the model accounted for stimulus duration and controlled for false positives when neural activation either scales or does not scale with stimulus duration. Our estimate is also unbiased, but contains high variance (Mumford et al., 2015) due to the degree of collinearity between stimulus duration and amplitude/velocity. This variability affects the sensitivity of univariate analysis, but the high sensitivity of MVPA can still identify relevant brain regions. In addition, our model assumed that neural activations in our study could potentially be related to stimulus duration at different levels for each condition, representing a full interaction model (Mumford et al., 2024). In a complementary analysis, we assessed whether assuming that neural activations relate to stimulus duration independent of condition would alter the trend of our results (Fig. S7). In this alternative model, increased variance prevented the inference of significant differences, though activation remained positively associated with amplitude and negatively associated with velocity (Fig. S7 A-D). A trend for higher activation during extension compared to flexion was also reproduced (Fig. S7E). Therefore, our full-interaction model effectively accounts for stimulus duration regardless of whether brain activation scales with it, and it retains power to infer significant differences across condition levels in univariate analysis.

Exps 1 and 2 used slow and rapid event-related designs, respectively. Rapid (compared to slow) event-related designs have a higher hemodynamic response functions (HRF), which can increase the variance of the estimate and consequently decrease sensitivity. On the other hand, rapid event-related designs are often preferred experimentally and analytically because they allow the inclusion of more trials and consequently more condition levels within a run. This was likely the reason why we observed idiosyncrasies between approaches only for this experiment for Exp 1. For example, amplitude was associated with S2 response in MVPA but not in univariate analysis and with thalamus and cerebellum in univariate but not in MVPA.

Finally, an alternative explanation for the negative association between activation and passive movement velocity may be due to the estimation procedure. Nurmi et al. (2018) reported that, in a block design, S1 activity gradually increased with movement rate, but plateaued or even decreased at higher rates. The authors observed that relatively lower rates better tracked the convolved HRF. Since both the univariate analysis and the MVPA in our study relied on the GLM, the results depend on how well the BOLD signal followed the convolved HRF. However, we argue that such a degradation of estimation may be less likely in an event-related design. Further studies may help clarify this point.

### 4.6. Limitations

The present study has several limitations. First, moving the index finger in opposition to the thumb is only one of a multitude of possible finger movements. Solutions in MRI-compatible robotics may use more degrees of freedom to explore and characterize the kinaesthetics of passive movement for other hand movements. Second, on a methodological level, only four runs were available for the cross-validation procedure to determine the performance of the predictive model, and a larger number of runs could have led to more robust results. Third, for training the model, we used a linear support vector machine (SVM) for the separation between classes, which might have led to relatively low accuracy values compared to non-linear models; however, the use of SVM improves interpretability and minimizes concerns about overfitting. Fourth, we used ROIs based on coordinates previously reported in related literature (Boscolo Galazzo et al., 2014).

Individualizing the definition of the ROIs, using functional or anatomical strategies, would presumably lead to more sensitive results for ROI analyses (Pamplona et al., 2020). Fifth, a previous study showed that MVPA classification performance is improved for rapid event-related designs through running separate GLMs for each trial and combining all the other trials into nuisance regressors (Mumford et al., 2012). The GLM approach we used, which consisted of estimating all the regressors of interest simultaneously, is a common procedure, but leads to higher estimation variability compared to separate GLMs. However, the same study concluded that estimating regressors of interest either separately or simultaneously results in similar classification performance when the design contains inter-stimulus intervals of at least 4 s on average, which was the case for Exp 1.

### 4.7. Conclusions

Here, we studied the association between brain regions and kinematic features of passive index finger movement, independently of each other. This was achieved through univariate and multivariate analyses, systematic robotically-driven passive movement stimulation, and controlling for stimulus duration, regardless of whether brain activation scales with it. We found that a large network of sensorimotor and motor regions encode the amplitude of passive finger movements, but neural correlates for velocity and direction of passive movement were weaker. The predominance of amplitude over other kinematic features suggests that brain activity related to passive movement is primarily related to the measurement of finger position through muscle spindles and to the constant updating of expected position. Understanding how passive movement is translated into neural responses is essential for designing neurally-guided passive movement therapy after brain injury. The findings presented here may help track neural reorganization and guide how neurorehabilitation can use neural representation of kinematic features to promote brain recovery.

## Acknowledgements

We thank Julio Dueñas for providing experimental data and Romy Lorenz for support with software.

## Ethics Approval Statement

This study used the data from a project approved by the local ethics committee (KEK 2010-0190).

## Data and Code Availability Statement

All obtained results and scripts used for the data analysis are available on the public GitHub repository: https://github.com/gustavopamplona/MVPA_passive_movement.

## Author Contributions

GSPP: conceptualization, methodology, software, validation, formal analysis, investigation, data curation, writing - original draft, writing - review & editing, visualization; JS: conceptualization, validation, writing - review & editing; EB: validation, writing - review & editing; OL: validation, writing - review & editing; SI: resources, writing - review & editing, funding acquisition; RG: conceptualization, validation, resources, writing - review & editing, funding acquisition; JLP: conceptualization, methodology, validation, writing - review & editing.

