## Supplementary material for "Neural correlates of kinematic features of passive finger movement revealed by univariate and multivariate fMRI analyses"

### Figures

#### Fig. S1

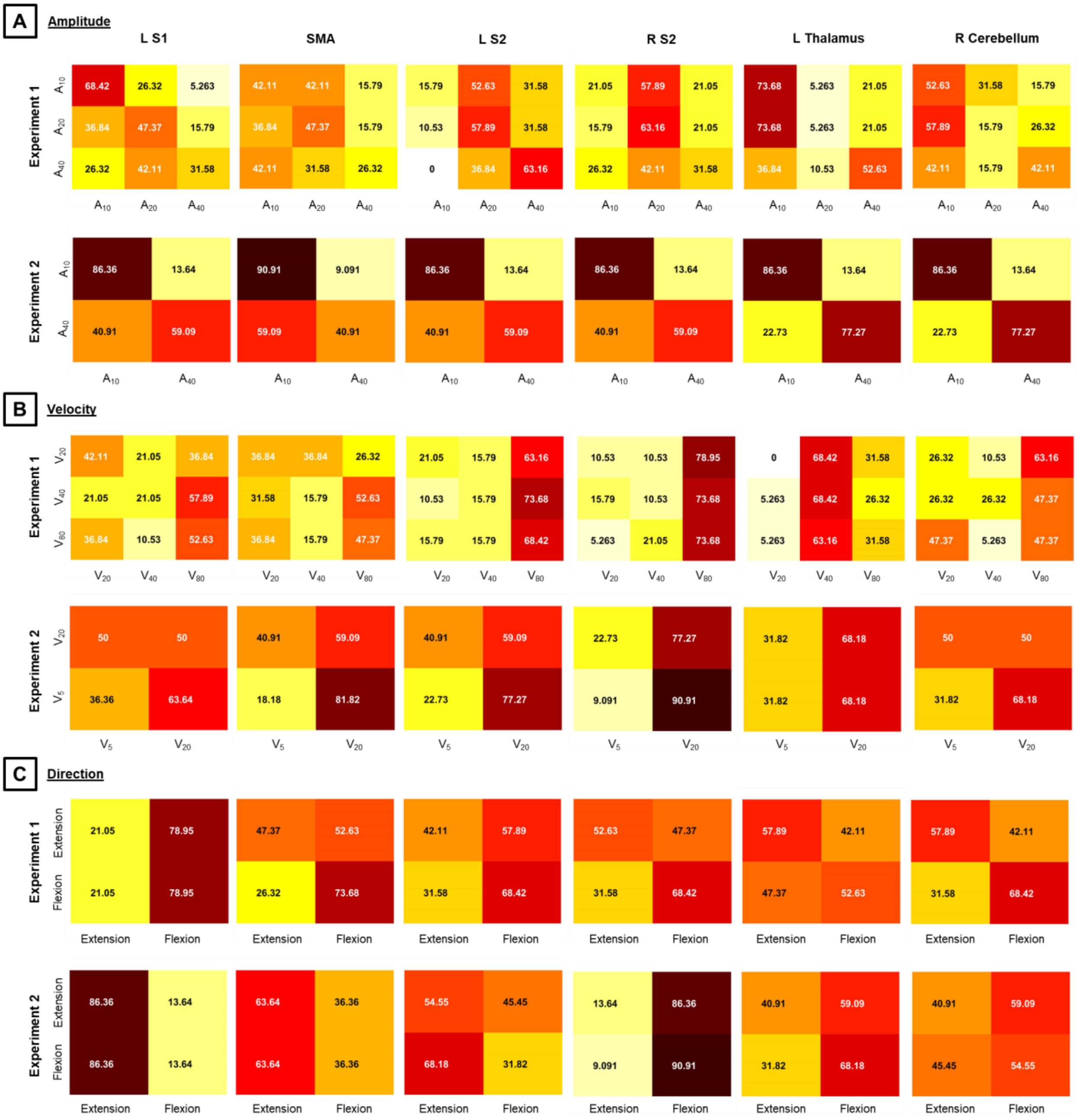

*Figure S1. Stability confusion matrices of MVPA results for (A) amplitude, (B) velocity, and (C) direction conditions and Experiments 1 and 2. Numbers in black (white) are higher (lower) than the chance level (33.33% for 3 conditions, 50% for 2 conditions). S1/S2 = primary/secondary somatosensory cortex, SMA = supplementary motor area, L/R = left/right.*

Fig. S2

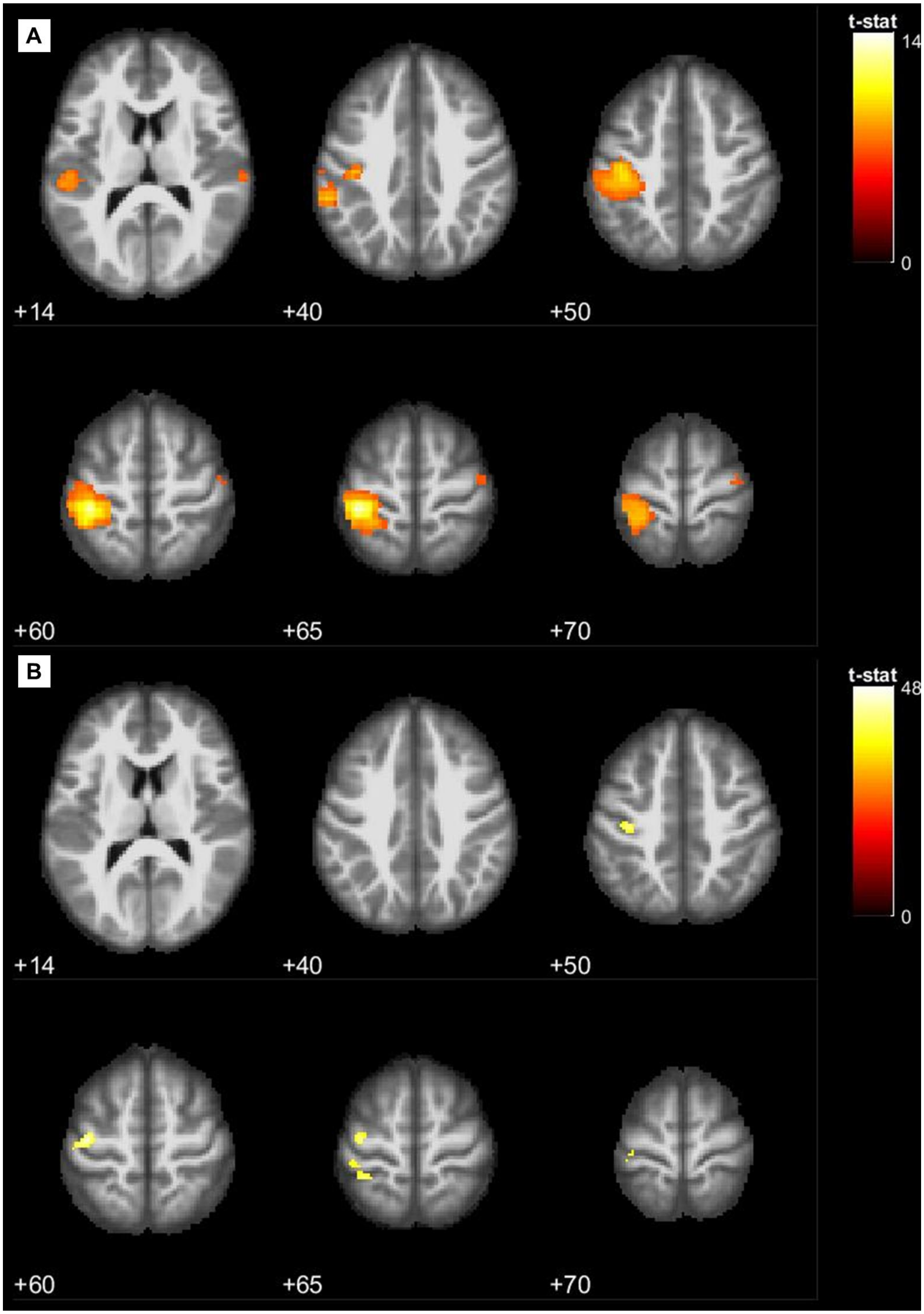


*Figure S2. Axial slices for whole-brain maps for amplitude in Exp 1 obtained with (A) MVPA and (B) univariate analysis.*

Fig. S3

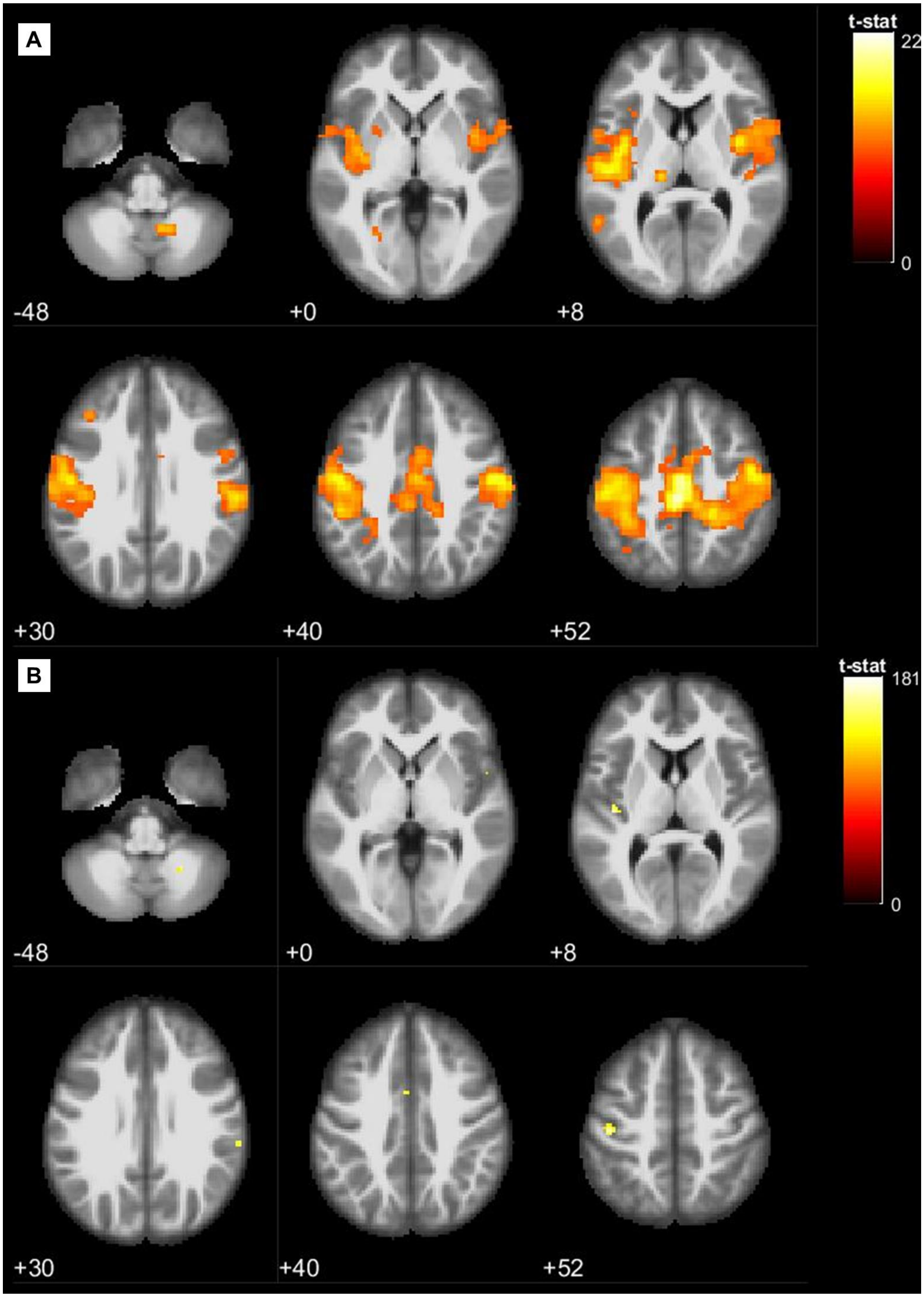


*Figure S3. Axial slices for whole-brain maps for amplitude in Exp 2 obtained with (A) MVPA and (B) univariate analysis.*

#### Fig. S4

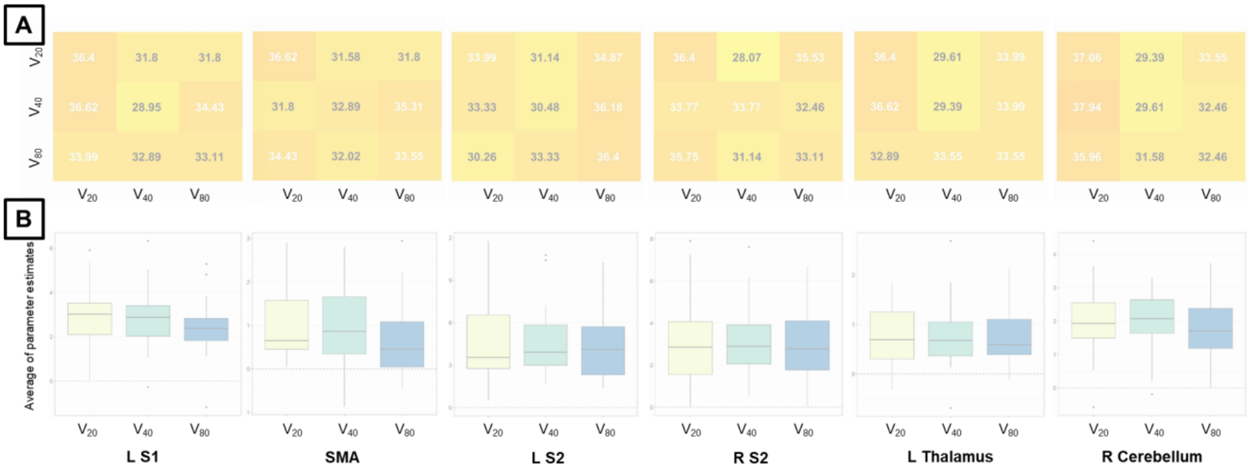


*Figure S4. MVPA and univariate analysis results for the velocity condition and Experiment 1. (A) ROI results for MVPA and (B) univariate analysis; in which the transparent plots represent non-significant discrimination for MVPA and $p > 0.05$ for the main effect of amplitude AND post-hoc pairwise differences for univariate analysis. Predictions in MVPA results were not significantly different from the chance level (33.33%), corrected for multiple comparisons using the Bonferroni method and bootstrapping (105 permutations) across subjects. There were no significant differences in univariate analysis results, corrected for multiple comparisons (Sidak method – ****: $p < 0.0001$). For MVPA confusion matrices, numbers in black (white) are higher (lower) than the chance level. S1/S2 = primary/secondary somatosensory cortex, SMA = supplementary motor area, L/R = left/right.*

Fig. S5

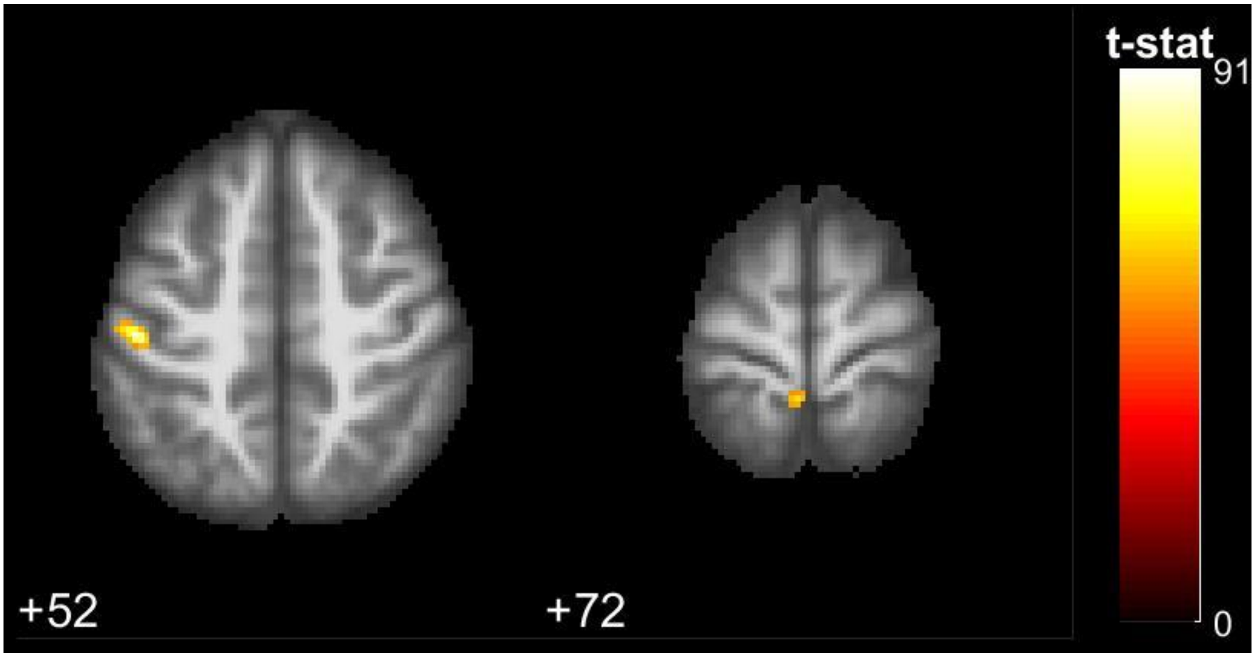


*Figure S5. Axial slices for whole-brain maps for velocity in Exp 2 obtained with univariate analysis.*

Fig. S6

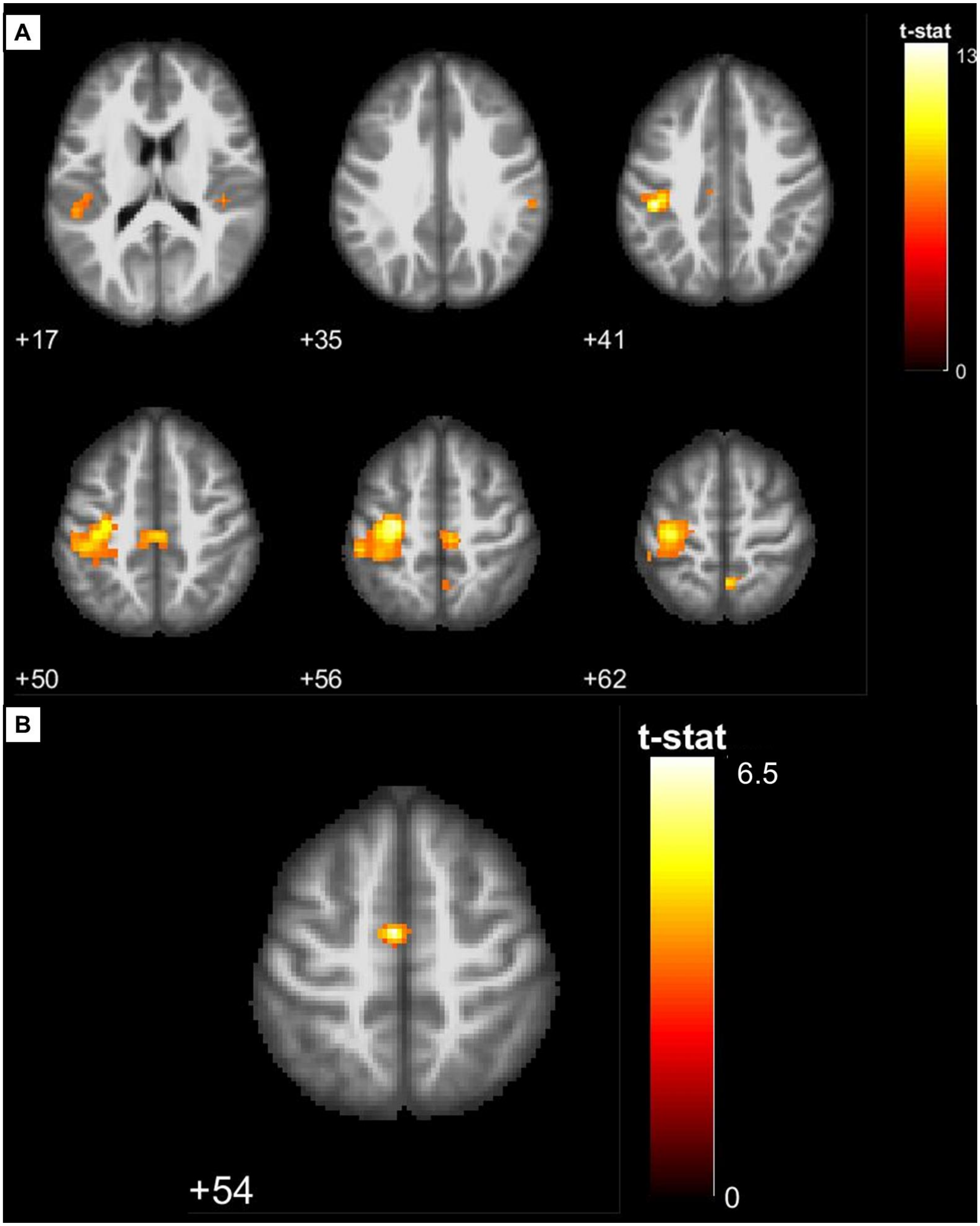


*Figure S6. Axial slices for whole-brain maps for direction in Exp 1 obtained with (A) MVPA and (B) univariate analysis.*

Fig. S7

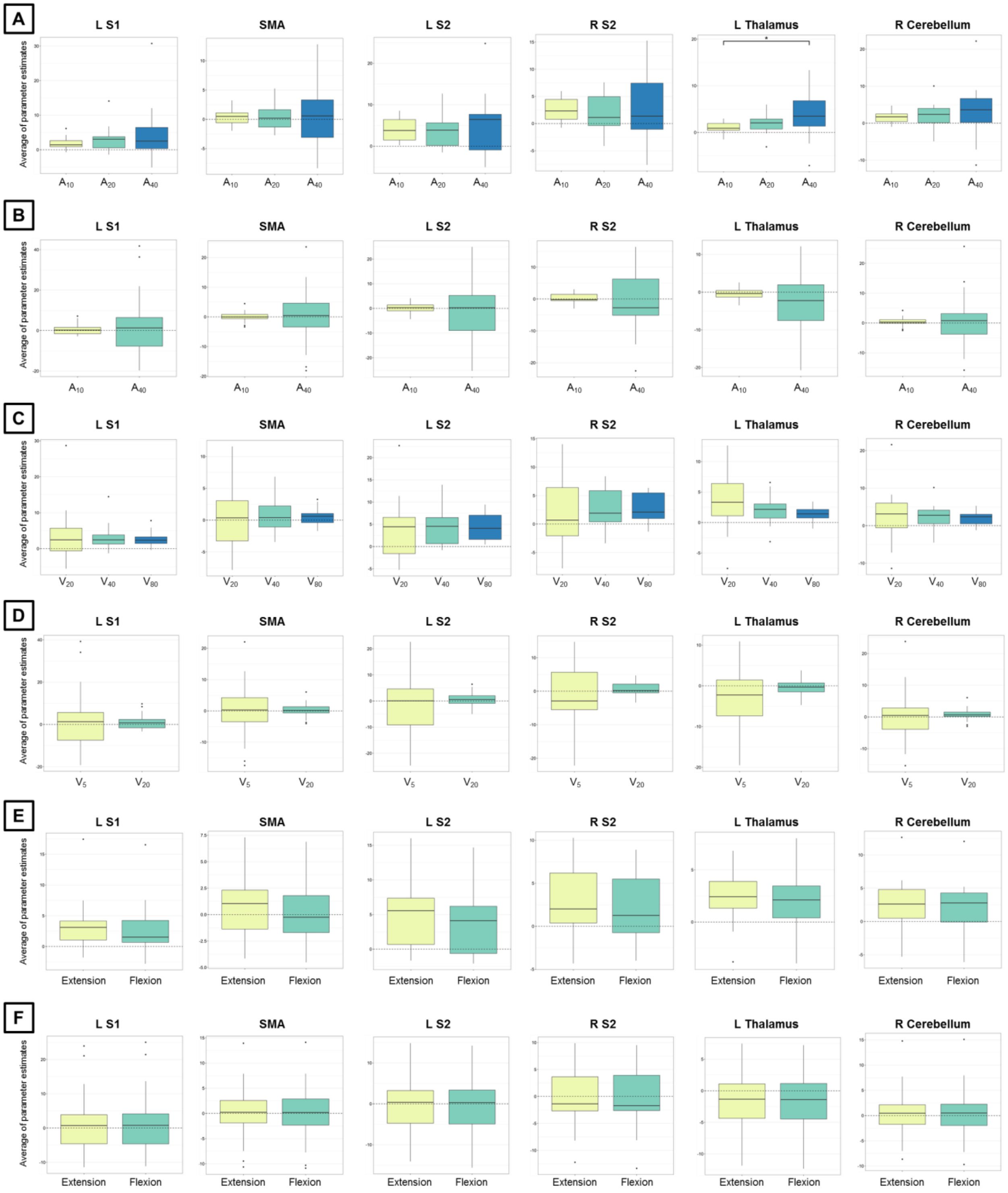


*Figure S7. Univariate ROI analysis results controlling for the interaction stimulus duration by kinematic feature. We show results for the amplitude condition and Exp 1 (A), amplitude condition and Exp 2 (B), velocity condition and Exp 1 (C), velocity condition and Exp 2 (D), direction condition and Exp 1 (E), and amplitude condition and Exp 2 (F). The*

*asterisk shows a significant post-hoc difference corrected for multiple comparisons (Sidak method, $p < 0.05$). S1/S2 = primary/secondary somatosensory cortex, SMA = supplementary motor area, L/R = left/right.*
